# Microbial cell factory optimisation using genome-wide host-pathway interaction screens

**DOI:** 10.1101/2023.08.30.555557

**Authors:** Paul Cachera, Nikolaj Can Kurt, Andreas Røpke, Tomas Strucko, Uffe H. Mortensen, Michael K. Jensen

**Author notes:** to whom correspondence should be addressed: Michael K. Jensen.

## Abstract

The ubiquity of genetic interactions in living cells challenges the concept of parts orthogonality, which is a cornerstone of synthetic biology. Parts, such as heterologously expressed genes, draw from shared pools of limited cellular resources and interactions between parts themselves and their host are inevitable. Instead of trying to eliminate or disregard these interactions, we propose to leverage them to promote desirable phenotypes. We recently described CRI-SPA, a method for high-throughput genome-wide gene delivery and screening of host:pathway interactions in *Saccharomyces cerevisiae*. In this study, we combine this method with biosensor-based high-throughput screening and high-density colony image analysis to identify lead engineering targets for optimising *cis-cis*-muconic acid (CCM) production in yeast cell factories. Using the biosensor screen, we phenotype >9,700 genotypes for their interaction with the heterologously expressed CCM biosynthesis pathway, including both gene knock-out and overexpression, and identify novel metabolic targets belonging to sulphur assimilation and methionine synthesis, as well as cellular redox homeostasis, positively impacting CCM biosynthesis by up to 280%. Our genome-wide exploration of host pathway interaction opens novel strategies for the metabolic engineering of yeast cell factories.

## Introduction

The field of synthetic biology aims at engineering biological functions *de novo*, either *in vitro* or *in vivo*, by cataloguing individual biological components as orthogonal parts (Canton et al., 2008). The notion that cellular systems can be packaged into independent modules is a useful abstraction, which facilitates studying biological complexity (Hartwell et al., 1999). This reductionist framework is enticing because it is conceptually simple and successfully rooted at the core of many engineering disciplines (Lazebnik, 2002). Bottom-up designs in living cells have been undeniably successful in the building of complex transcriptional logics (Friedland et al., 2009; Nielsen et al., 2016; Zong et al., 2017), and orthogonal transcription and translation of non-canonical amino acids (d’Aquino et al., 2018). However, such successes are built upon a thorough understanding of the underlying molecular processes and this understanding is lacking for numerous cellular mechanisms even in model organisms (Wood et al., 2019)

Reducing the complexity of interactions between heterologous gene products and their host to a handful of inputs-outputs is likely an oversimplification. Indeed, heterologously produced enzymes might interfere with the host cellular processes by e.g. exhausting pools of cofactors and energy, catalysing side reactions and producing toxic intermediates (Boo et al., 2019; Dahl et al., 2013; Sun et al., 2017; Wu et al., 2016). This challenge escalates dramatically when the number of coding sequences and edits that current metabolic engineering efforts currently introduce into cell factories is considered (Zhang et al., 2022).

The concept of host-aware synthetic biology has gained momentum in the past 10 years with a number of efforts addressing interactions between the synthetic system and the host (Boo et al., 2019; Cardinale and Arkin, 2012; de Lorenzo, 2011). For example, studies have quantified and attempted to reduce the heterologous burden of synthetic systems on the host (Borkowski et al., 2018; Ceroni et al., 2015; Gyorgy et al., 2015; Shopera et al., 2017; Weiße et al., 2015). An alternative approach is to reduce the cross talk between parts as well as between parts and the host, a strategy known as ‘isolation’ (Darlington et al., 2018; Lou et al., 2012; Meyer et al., 2015; Segall-Shapiro et al., 2014). A burgeoning branch of synthetic biology goes as far as proposing to fully isolate human-made mechanisms through a complete orthogonal dogma (Costello and Badran, 2021; Liu et al., 2018).

Instead of eliminating genetic interactions, we reason that they may serve an engineering task. Indeed, the systematic combination of double KO in yeast revealed that, in rare instances (≈1.5%), the interaction of two gene KOs could yield an unexpected increase in fitness (Costanzo et al., 2016, 2010). This hints that sporadic phenotypic maximas hidden in the interaction landscape are awaiting to be identified and exploited. This idea is supported by adaptive laboratory evolution and multiplexed CRISPR-based screens which find that combinations of edits often act synergistically to yield a sought-for phenotype which cannot be reduced to the sum of its components (Bao et al., 2018; Lian et al., 2017; Wang et al., 2020).

The identification of rare phenotype maxima in the interaction landscape requires the means to generate and analyse strain variants at high throughput. We recently described CRI-SPA, a method capable of delivering a genetic feature to an arrayed yeast library within a week (Cachera et al., 2023). In its first version, we used CRI-SPA to screen the yeast knowck-out (YKO) library for gene knockouts interacting with the biosynthesis of the plant pigment betaxanthin. The workflow made use of the yellow phenotype produced by betaxanthin to extract a productivity readout with image analysis. Image analysis is arguably the simplest and most inexpensive high-throughput readout strategy. Unfortunately, only a minority of value-added products, such as betaxanthin and carotenoids, directly produce a visible phenotype (Zeng et al., 2020). Biosensors, which translate a chemical input into a measurable output, typically fluorescence, widen the range of products and cellular processes which can be measured and quantified (d’Oelsnitz et al., 2022; Rogers et al., 2016). A biosensor can be expressed in the producing strain and scale to any throughput with no additional cost. In addition, a wide collection of biosensors have been developed over the past decades, opening their application to a large variety of products (Koch et al., 2019).

Here, we couple CRI-SPA to the biosensing of the otherwise inconspicuous bio-plastic platform chemical *cis-cis*-muconic acid (CCM) with the aim of identifying novel host:pathway interactions in *S. cerevisiae*. Since CCM was first synthesised in yeast a decade ago (Curran et al., 2013b), numerous metabolic engineering efforts have attempted to increase its production by improving the availability of shikimate pathway precursors or by relieving the enzymatic bottleneck of the rate-limiting conversion of protocatechuic acid (PCA) to catechol (**Fig. 1a**), (Brückner et al., 2018; Curran et al., 2013a; Suástegui et al., 2017; Weber et al., 2012, 2017). CCM is the ideal test-bed for our study because identification of rational engineering strategies to increase its titers likely have reached a plateau. On the other hand, the engineering of biosensors for CCM (Skjoedt et al., 2016; Snoek et al., 2020) and for shikimate pathway end products (Leavitt et al., 2016) have supported a number of screens and directed evolution studies (Jensen et al., 2021; Leavitt et al., 2017; Snoek et al., 2018; Wang et al., 2021, 2020). Biosensor-coupled directed evolution can efficiently isolate strains with higher production capacity. However, disentangling the mechanisms behind these improvements and their reverse engineering can be complicated by the number of mutations which need to be tested individually or in combination (Mundhada et al., 2017; Wang et al., 2020).

**Fig. 1.**
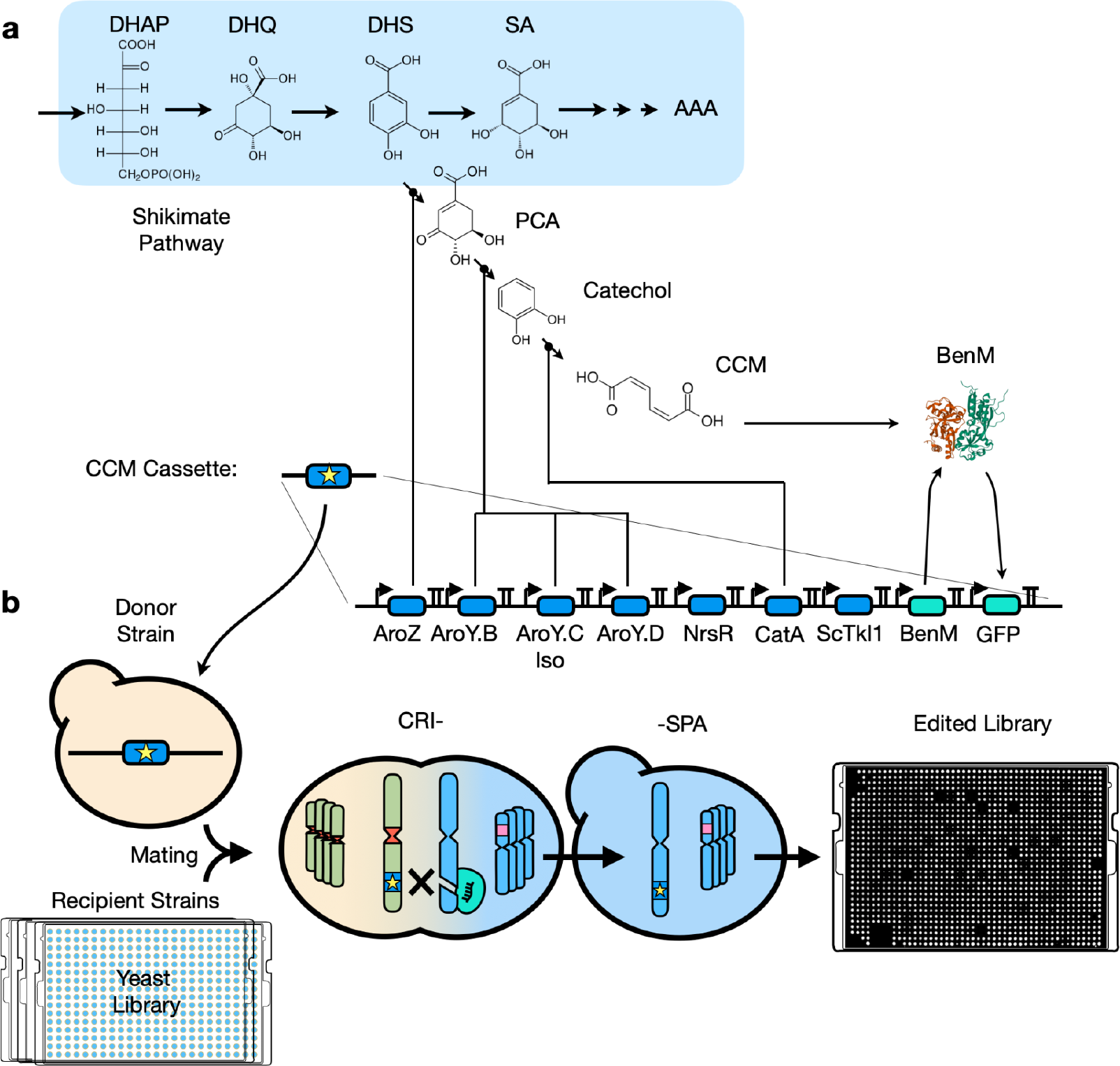
Overview of the design of the CRI-SPA screen for identification of host genes interacting with CCM synthesis. **a**) Schematic of the synthesis and reporting of CCM by the CCM cassette. Abbreviations: AroZ: *Klebsiella pneumoniae* DHS dehydratase. AroY.B-AroY.Ciso-AroyD: *K. pneumoniae* PCA decarboxylase cluster (iso=isoform). NrsR: Nourseothricin Resistance, CatA: *Acinetobacter baylyi* catechol 1,2-dioxygenase DAHP: 3-Deoxy-D-arabinoheptulosonate 7-phosphate, ScTKL1: *S. cerevisiae* Transketolase, BenM: engineered CCM biosensor. DHQ: 3-Dehydroquinic acid, DHS: 3-dehydroshikimate, SA: shikimic acid, AAA: Aromatic Amino Acids. Refer to **Suppl. Fig. S1** for all parts in the cassette and **Suppl. Table S1** for the sourcing of these parts. **b**) The CCM cassette is inserted in the CRI-SPA donor strain and delivered to all strains of a yeast library by CRI-SPA. Final colony fluorescence can be extracted with image analysis as a proxy for quantifying CCM titers.

To open new engineering avenues for CCM synthesis, we sought to screen for beneficial host:pathway interactions resulting in an increased CCM accumulation in *S. cerevisiae*. We used CRI-SPA to deliver a cassette encoding for the synthesis and reporting of CCM to the strains of the genome-wide OverExpression (OEx) and Knock-Out (KO) libraries. We measured CCM biosensing directly from colonies growing on agar plates enabling high-density array image analysis and identified known and novel cellular functions interacting with CCM synthesis. Importantly, many of the positive hits identified in the screen were subsequently validated and shown to outperform the reference strain in liquid medium.

## Results

### Biosensor-assisted CCM quantification based on image analysis

To enable a genetically-encoded biosensor to report a proxy for CCM formation, we engineered the CRI-SPA donor strain (Cachera et al., 2023) with an expression cassette encoding the CCM biosynthesis pathway and the CCM biosensor (**Fig. 1a; Supplementary Figure S1**). Engineering of the CRI-SPA donor strain was initiated by building a single 19kb DNA cassette able to both synthesise and report CCM (**Fig. 1a**). Subsequently, the CCM cassette is then designed to transfer from the donor strain to yeast libraries using CRI-SPA, (Giaever et al., 2002; Weill et al., 2018; Yofe et al., 2016), and edited strains of the library screened for fluorescent signals as a proxy for CCM synthesis (**Fig. 1b**).

Specifically, for CCM synthesis, we refactored the previously described biosynthetic pathway (Skjoedt et al., 2016; Snoek et al., 2020) by replacing all repetitive promoters and terminators with characterised strong equivalents (Curran et al., 2013a; Zhang et al., 2020) (**Suppl. Figure S1**). This minimises the risk of repeat-recombination events within the pathway that may result in gene rearrangement or gene copy loss during regular strain propagation or during CRI-SPA-mediated cassette transfer. We also engineered an additional donor strain expressing the decarboxylase AroY-B variant (AroY-B_P146T), which we previously evolved for optimized conversion of PCA to catechol (Jensen et al., 2021, **Fig. 2a**). For CCM detection, an improved version of the BenM biosensor (BenM17_D08, henceforth referred to as BenM) was encoded in the 19 kb cassette (Snoek et al., 2020).

**Fig. 2.**
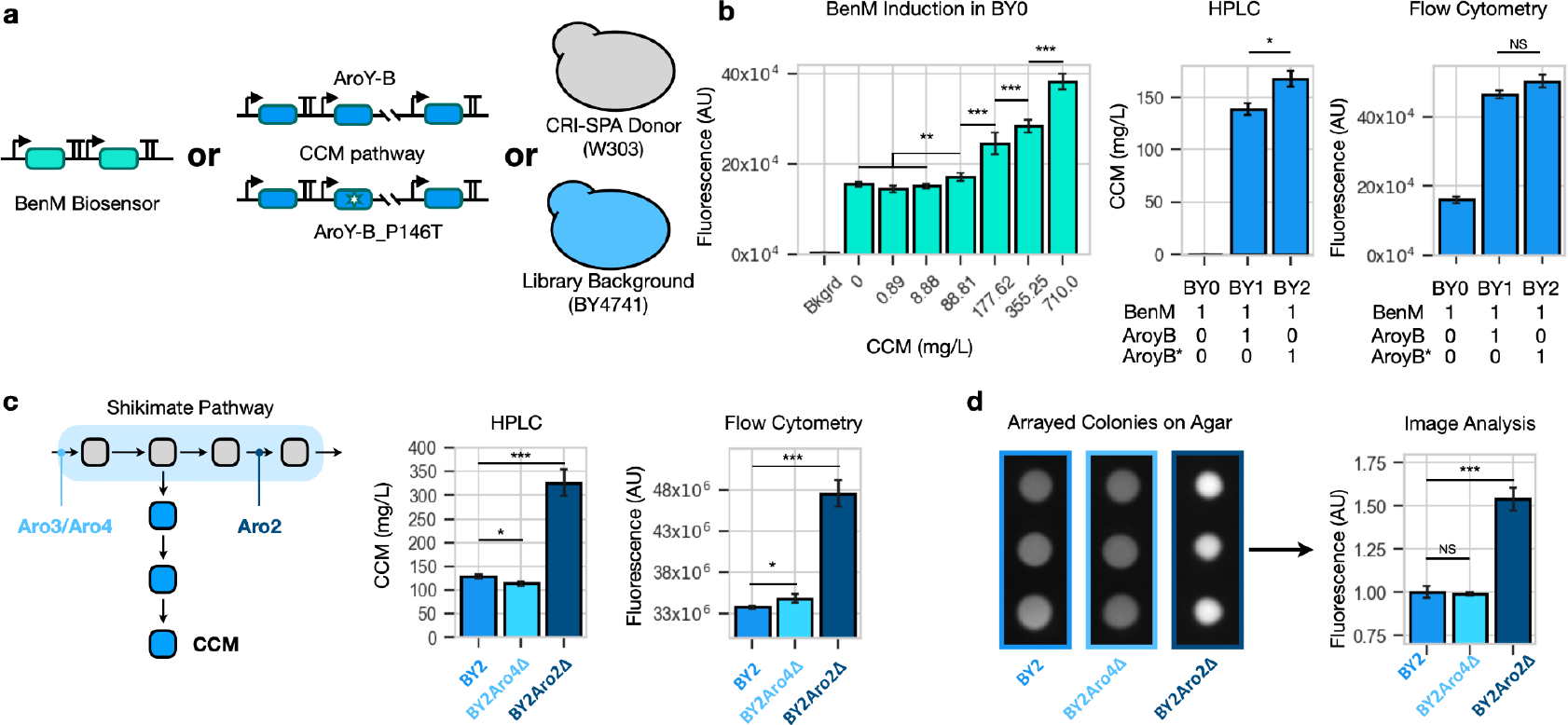
Characterization of CCM production and biosensing in CRI-SPA engineered yeast. **a**) The two variants of the CCM cassette, harbouring either the WT AroY-B decarboxylase or the optimised variant AroY-B_P146T, were introduced in the CRI-SPA donor strain and the BY4741 background. **b**) Left: Induction of the CCM biosensor in strain BY0. Centre: CCM titers for strains BY0, BY1 and BY2. Right: fluorescence measured by flow cytometry for strains BY0, BY1 and BY2. **c**) KO of genes *ARO2* and *ARO4* as positive and negative controls, respectively. Left: the reactions catalysed by Aro2 and Aro4 are upstream and downstream of the CCM branch. Centre: CCM synthesis by *aro2*Δ and *aro4*Δ KO strains. Right: Fluorescence of *aro2*Δ and *aro4*Δ strains. **d**) Changes in CCM titers cause a fluorescent change quantifiable directly from agar colonies. Fluorescence is expressed in fold changes relative to BY2. For b) and d), bars show the mean of at least three experimental replicates. For b) and c), replicates are technical. Biological replicates of the experiments can be found **Suppl. Fig. S2**. For all graphs, significance was assessed with a two-sided t-test *<0.05, **<0.01, ***<0.001. AU: arbitrary units.

For CRI-SPA-mediated editing, we inserted the CCM cassettes in the XII-5 donor site of the CRI-SPA donor strain of both mating types (Cachera et al., 2023). For the *Mat-****a*** donor strains, this resulted in strain CRI-SPA Donor strain CD1 (*Mat-****a*** with expression of WT *AroY-B*) and CD2 (*Mat-****a*** with optimised AroY-B_P146T). For the *Mat****-alpha*** donor strains, the cassettes were chromosomally inserted and the expression cassette encoding nourseothricin resistance was switched to kanamycin resistance, resulting in CD3 (*Mat-alpha* with WT AroY-B) and CD4 (*Mat-alpha* with optimised AroY-B_P146T). In parallel to these CDs, we built two control strains, BY1 and BY2 harbouring the two pathway variants with WT AroY-B or the optimized AroY-B_P146T in background BY4741. BY4741 is commonly used in the making of yeast libraries (Giaever and Nislow, 2014; Weill et al., 2018; Yofe et al., 2016)(**Fig. 2a**). BY1 and BY2 thus serve as controls identical to the final edited library that the CRI-SPA screens will produce. Finally, to test the inducibility of BenM in the library background, we also built BY0 a strain harbouring the CCM biosensor but without the CCM biosynthetic pathway.

Next, we validated the production of CCM and its biosensing in the donor strain and the library controls, and found that the refactored pathways produced CCM at levels in agreement with previous work (Jensen et al., 2021)(**Fig. 2b**). We also detected the expected titer increase caused by the optimised AroY.B decarboxylase (WT: 138.7 mg/L, optimised: 168.0 mg/L, two-sided t-test, P-value = 0.012). Furthermore, fluorescence analysis measuring production of the GFP reporter confirmed the inducibility of the biosensor by CCM, albeit with a lower dynamic output range (2.5x) than previously reported (Snoek et al., 2020)(**Fig. 2b**). Encouragingly, the biosensor was able to detect the modest, yet insignificant, increase in CCM production caused by the optimised AroY.B compared to the WT decarboxylase, indicating that the level of CCM produced by the CCM biosynthetic pathway fell within the operational range of detection of the biosensor (**Fig. 2b**).

To further test the ability of the biosensor to detect differences in CCM biosynthesis caused by gene edits in CRI-SPA strains, we built additional control strains that are expected to show increased or decreased CCM synthesis. Here, we deleted *ARO4* encoding DAHP synthase and *ARO2* encoding chorismate synthase, which act up- and downstream of the branchpoint to the heterologous CCM pathway, respectively (Jones et al., 1991; Künzler et al., 1992). The genes were deleted in BY2, the most productive control strain variant, resulting in strains BY2Aro4Δ and BY2Aro2Δ which were cultivated in liquid Synthetic Complete (SC) medium for 72 h (**Fig. 2c**). For the *aro2*Δ strain, we observed more than 2-fold increase (p-value < 0.001) in CCM titers. The *aro4*Δ strain displayed only a modest reduction in CCM production relative to the wild-type strain (WT: 129.9 mg/L, *aro4*Δ: 113.5 mg/L, p = 0.034). This is probably due to the activity of redundant *ARO3* which can maintain the flux through the shikimate pathway even in the absence of *ARO4* (Curran et al., 2013b). Importantly, quantification of the GFP reporter by flow cytometry of liquid cultures of strains BY2, BY2Aro4Δ and BY2Aro2Δ correlated with the CCM levels obtained by HPLC, confirming that the biosensor was able to report changes in CCM synthesis caused by single gene deletions in CRI-SPA strains (**Fig. 2c**).

To match the high-density agar array format of CRI-SPA, we investigated the possibility of reporting CCM productivity from BY2, BY2Aro4Δ and BY2Aro2Δ directly from colonies growing on agar. In this case, image analysis of colonies of strain *aro2*Δ detected a 56% increase (p-value < 0.001) in colony fluorescence intensity compared to the reference strain (**Fig. 2d**). Importantly, the fluorescence intensity observed for BY2, BY2Aro4Δ and BY2Aro2Δ again correlated with the analytical quantification performed by HPLC (**Fig. 2c**).

In summary, we demonstrated that when our CCM biosensor is implemented in a setup mimicking CRI-SPA output strains, it is able to differentiate host KOs causing an increase in CCM synthesis under screening conditions.

### High-density image analysis of >9,700 genotypes

Next, we sought to employ this framework to screen for genome-wide host:pathway interactions for all genes present in the YKO and OEx libraries. For this purpose, we employed CD2 to deliver the CCM cassette to all strains of the YKO and OEx libraries, interrogating fluorescence signals as a proxy for CCM accumulation in a total of >9,700 arrayed mutant strains. As expected, the arrayed colonies edited by CRI-SPA were fluorescent, hinting to a successful transfer of the 19kb CCM cassette into the libraries (**Suppl. Fig. S3a**). We also observed that positional artefacts, which are known to affect colony size in high-density arrays, also affected the colony fluorescence in our screen (Baryshnikova et al., 2010, **Suppl. Fig. S3b-c**). To mitigate this phenomenon, we applied the data processing methods developed by the Boone and Andrews labs to correct for positional artefacts and to remove colony outliers from our data (Wagih et al., 2013).

Following this processing step, we ranked the genes in the two libraries according to their corrected fluorescence. Here, we found that the screen of the KO library had a stronger signal-to-noise ratio than the OEx screen (**Fig. 3a, Suppl. Fig. S3a**). The *Z* scores for the first and last 2% quantiles were 9.45 and -15.64 for the KO screen, respectively, but only 4.21 and -4.83 for the OEx screen. This difference in dynamic range was also qualitatively visible on raw screen pictures, where the CRI-SPA colonies on the KO library plates displayed more variations in fluorescence intensity compared to the CRI-SPA colonies on the OEx library plates (**Suppl. Fig. S3a**). Yet, and most importantly, the fluorescence signal was visible on quadruplicate colony images recovered from the images of both screens (**Fig. 3a-b**). Furthermore, repeating the YKO library screen, we showed that the fluorescent signal obtained from the CRI-SPA colonies were reproducible (PCC = 0.71, p<0.001, **Fig. 3a, insert**). Combining the data produced by the two YKO screen repeats and the single OEx screen (see materials and methods), our final dataset quantified the effects of 4,596 gene KOs (97.8% of the YKO library) and of 4,983 genes overexpressed (99.0% of the OEx library) on the synthesis of CCM.

**Fig. 3.**
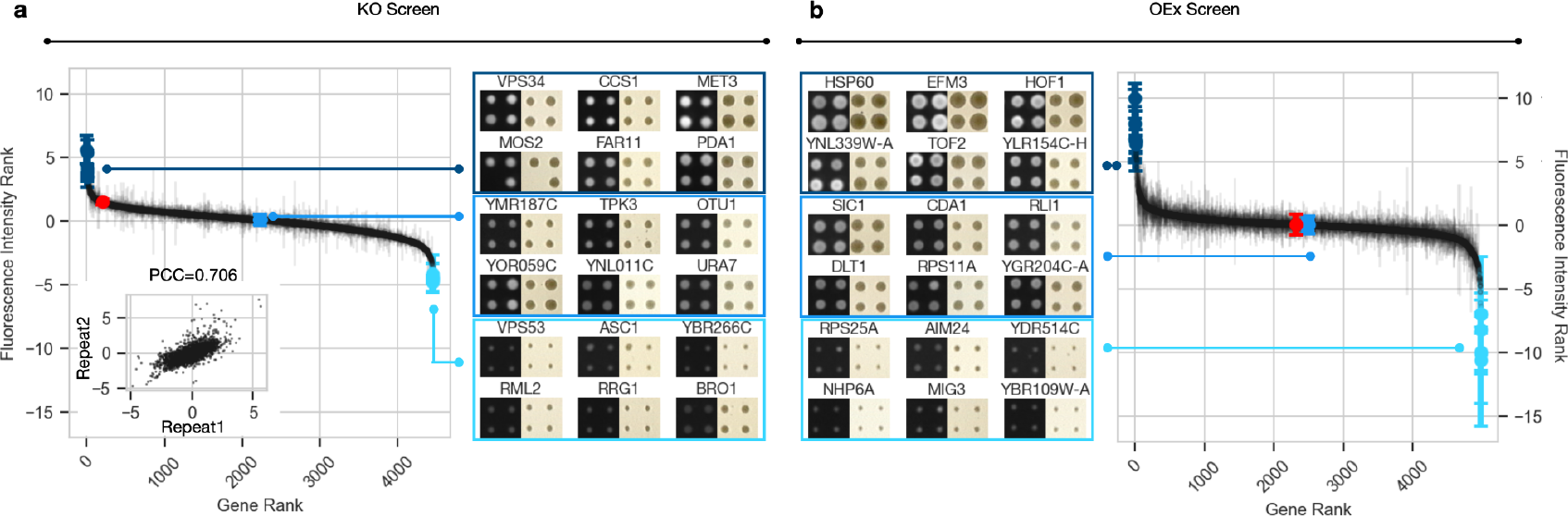
Ranking of host:pathway interactions for gene KO and overexpression. **a**) Left: Ranking the fluorescence intensity scores for 4,452 gene KO obtained with the first repeat of the screen on the YKO library. Inset shows reproducibility of the fluorescence intensity scores obtained for each mutant strain in the gene YKO library between two independent trials. Right: Fluorescent and non-fluorescent images of colonies from the same plate sampled at the indicated ranks. **b**) Right: Ranking the fluorescent intensity scores for 4,983 gene OEx strains obtained with the OEx strain library. Left: Fluorescent and non-fluorescent images of colonies sampled at the indicated ranks. Each point is the mean fluorescence of three or more independent CRI-SPA colonies, the error bar is the standard deviation. Red points indicate the WT library strain edited by CRI-SPA.

### CCM biosensor identifies known host:pathway interactions

Encouragely, we found that the ranking produced by the KO and OEx fluorescence datasets captured the beneficial effects of knocking-out *ZWF1* and overexpressing of *PAD1*, two common targets for CCM overproduction in *S. cerevisiae* (Curran et al., 2013b; Weber et *al*., *2012*). Indeed, deletion of *ZWF1*, which encodes a cytoplasmic glucose-6-phosphate dehydrogenase committed to the first step of the pentose phosphate pathway, ranked high in fluorescence at position 358 out of 4,596 in the KO screen (**Fig. 4, Suppl. Table S2**). Deletion of *ZWF1* in combination with overexpression of *TKL1*, encoding transketolase 1 responsible for the conversion of xylulose-5-phosphate and ribose-5-phosphate to sedoheptulose-7-phosphate and glyceraldehyde-3-phosphate, is a modification recurrently employed to rewire the flux of the pentose phosphate pathway (PPP) towards its non-oxidative branch (Brückner et al., 2018; Curran et al., 2013b). This increases the availability of E4P, a precursor known to be rate limiting for the shikimate pathway and CCM production (Horwitz et al., 2015; Suástegui et al., 2016). Since *TKL1* is encoded in the CCM cassette delivered to the libraries **(Fig. 1a)**, the high rank of the *ZWF1* deletion strain corroborates previous strategies for improving CCM synthesis.

**Fig. 4.**
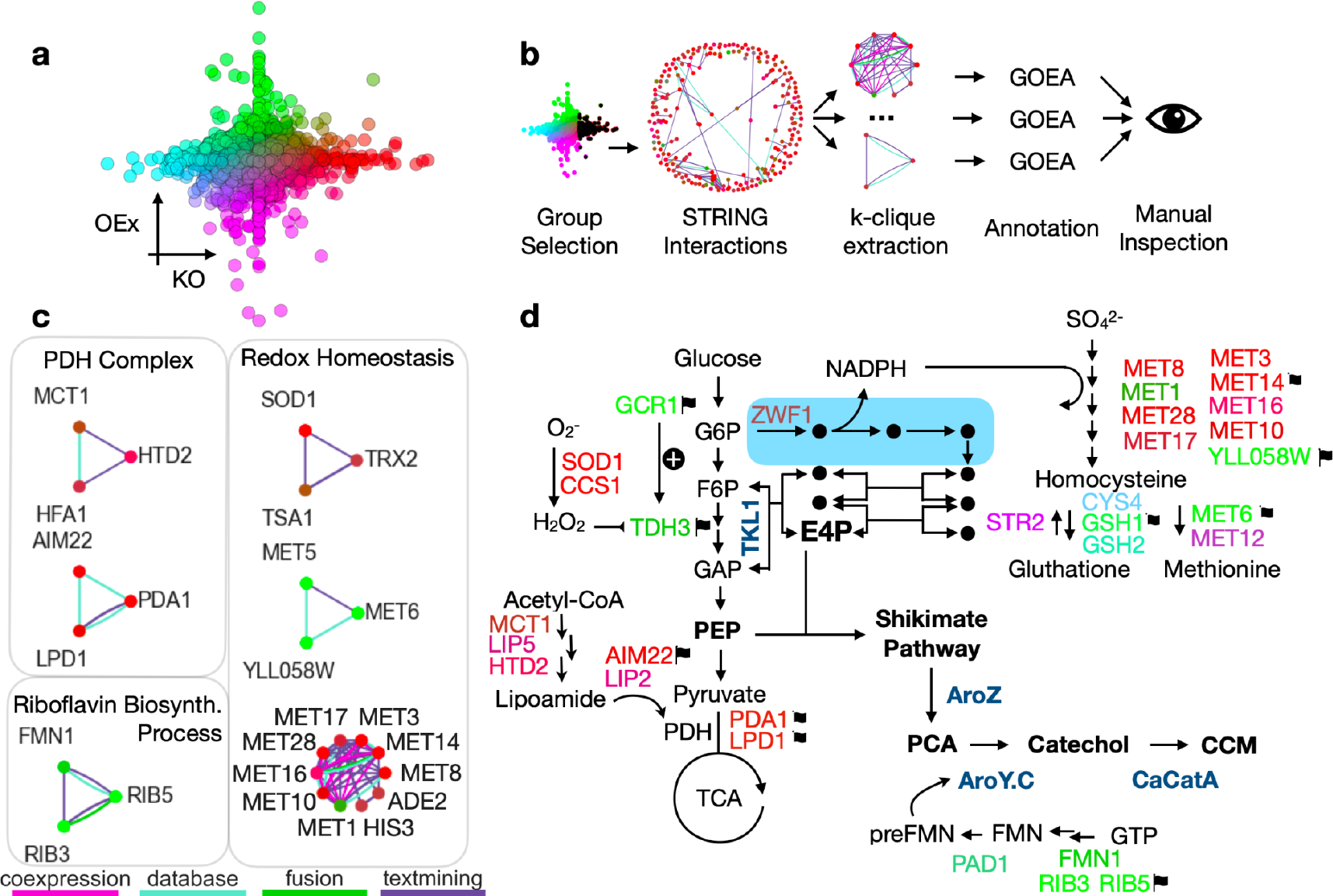
New cellular functions interacting with the synthesis of CCM. **a)** KO-OEx colour map. Each point is the mean fluorescence of a gene in the KO screen (x-axis) and the OEx screen (y-axis). Each gene was colour-coded according to its x-y coordinate to map the effects of its KO and OEx on CCM synthesis as shown in panels b-d. **b)** Overview of the hit gene cliques extraction and annotation workflow. Group selection: a group of hits is selected from the screen data. For e.g. KOs causing the strongest fluorescence (black dots). STRING interactions: A STRING interaction network is drawn for the selected group. k-clique extraction: cliques of genes linked by STRING interactions are isolated with the k-clique extraction algorithm. Annotation: GOEA is performed for the set of genes present in the extracted cliques to find enriched GO terms. Manual inspection: the enriched GO terms are inspected for cellular functions interacting with CCM synthesis. **c)** Selected cliques extracted in b) showing mechanistically interpretable interactions with CCM synthesis. Node colours are determined by the colour map in a); the type of STRING interaction is indicated by the bottom legend. **d)** Mapping of the relationship of gene hits found in cliques and discussed in text with CCM synthesis. Gene name colours are determined by colour map in a), genes selected for HPLC validation are indicated with a flag, genes encoded by the CCM cassette are shown in bold blue. CCM intermediates and precursors are shown in bold black.

Likewise, we found *PAD1*, encoding phenylacrylic acid decarboxylase 1, to rank at position 231 out of 4,983 in the OEx screen (**Fig. 4, Suppl. Table S2**). Pad1 catalyses the prenylation of riboflavin, that is the same reaction catalysed by the cassette-encoded AroY-B (Weber et al., 2017). Prenylated riboflavin is a necessary cofactor in the decarboxylation of PCA to catechol catalysed by AroY.Ciso **(Fig. 1a)**. AroY.Ciso has been shown to be rate limiting in the synthesis of CCM in yeast (Brückner et al., 2018; Curran et al., 2013b; Leavitt et al., 2017; Payer et al., 2017; Weber et al., 2012). The prenylation catalysed by Pad1 is slow (k_cat_ = 12.2 h^-1^), yet more effective than AroY-B, and co-expression of Pad1 with AroY.C is known to improve availability of prenylated riboflavin (Arunrattanamook and Marsh, 2018; Weber et al., 2017). Altogether, the high rank of *PAD1* in the OEx screen supports the current understanding that insufficient prenylated riboflavin is responsible for the AroY.C bottleneck in CCM synthesis.

### Network analysis identifies novel metabolic engineering targets

We next sought to obtain a broader system-level knowledge of our data in search of novel cellular functions interacting with the synthesis of CCM. We envisioned that gene:gene interactions could be employed to identify clusters of functionally related genes interacting with CCM synthesis. To visualise these interactions, we built networks for the 250 highest- and lowest-ranking gene groups from the YKO and OEx library screens based on associations stored in the STRING database (Szklarczyk et al., 2021)(**Suppl. Fig. S4**). In each of the 250 hit groups, we observed a subset of genes forming densely connected clusters, mostly through ‘database’ and ‘coexpression’ interactions, among a larger number of unconnected genes (**Suppl. Fig. S4**). This suggested that functional gene clusters were indeed present in each hit group.

We isolated these with a k-cliques detection algorithm (Palla et al., 2005). This approach is used to find all cliques (i.e. complete subgraphs) connected to each other by k-1 connections. We set k=3 and a threshold of number of nodes ≥3 to identify cliques with at least three nodes in which all nodes were connected to the rest of the clique with at least k-1=2 interactions. Next, we labelled these cliques, by running Gene Ontology Enrichment Analysis (GOEA) for the gene sets included in them. As a control we also ran this analysis for a group made of 250 randomly selected genes across the dataset.

With this workflow (**Fig. 4a-b**) we found that all hit groups produced at least ten cliques (**Suppl. Fig. S5**). Besides the lowest-ranking 250 group from the YKO library screen, which produced nine cliques with five or more genes, most cliques in all other groups had only three genes. GOEA labelling of the cliques indicated that they represented a wide variety of cellular functions, including multivesicular body sorting and rRNA processing (**Suppl. Fig. S5**). We also note that the control group of randomly chosen genes also produced cliques with GOEA labels (**Suppl. Fig. S5e**). This advised caution as cliques, i.e. networks of functionally related genes, might simply be found by chance when randomly drawing a set of 250 genes from the *S. cerevisiae* gene pool. We therefore carefully inspected the cliques in the hit groups in search of mechanistically meaningful interactions with CCM synthesis.

Firstly, we noticed the presence of a clique enriching the GO term “riboflavin biosynthetic process” composed of genes encoding Fmn1, Rib5 and Rib3, all taking part in riboflavin synthesis (**Fig. 4c**). As mentioned earlier, the high ranking of *PAD1* in the OEx screen, suggests that the supply of the prenylated riboflavin cofactor is limiting. Here, the high-ranking of *FMN1, RIB5* and *RIB3* OEx hits suggests that, in addition to its prenylation, insufficient supply of riboflavin might also contribute to the AroY.C decarboxylation bottleneck (**Fig. 4d**).

Secondly, we found three cliques involved in cellular redox homeostasis. This included two cliques taking part in the process of sulphur assimilation for *de novo* methionine synthesis which is the cell’s most important electron sink (Rashida and Laxman, 2021; Thomas and Surdin-Kerjan, 1997). Interestingly, one of these two cliques was found in the top KO group (containing *MET1,3,8,10,14,16,17,28, HIS3* and *ADE*2) while the other in the top-ranking OEx group (containing *MET5,6* and *YILL058W*, **Fig. 4c**). The third clique, linking *SOD1, TRX2* and *TSA1* was labelled with ‘antioxidant activity’, and was extracted from the top-ranking hits of the KO screen. The Sod1-chaperone Ccs1, although not included in the clique, was also the strongest hit in the KO screen (**Suppl. Table S2)**. Last, we also noticed that the synthesis and recycling of glutathione, directly downstream of methionine synthesis, formed a hit hotspot, although it didn’t result in the identification of a clique.

Although less direct, the interaction between redox homeostasis and the CCM pathway becomes clear when recognising that E4P availability, CCM’s rate limiting precursor (Suástegui et al., 2016), increases upon shutdown of the oxidative branch of the PPP (Curran et al., 2013b). The oxidative PPP is the cell’s most important source of NADPH and its up-regulation is critical to maintain the NADPH/NADP ratio (Montllor-Albalate et al., 2022; Ralser et al., 2007). Hence, decreased NADPH consumption in mutants unable to assimilate sulphur should inhibit the oxidative PPP and improve E4P availability. Conspicuously, a recent study proposed that Sod1, a superoxide dismutase, had a crucial role in upregulating the oxidative PPP in response to oxidative stress. This study was found that under oxidative stress, conversion of superoxide into H_2_O_2_ by Sod1 directly inhibits the *TDH3-*encoded glycolytic enzyme Gapdh which redirects carbon flux into the oxidative PPP (Montllor-Albalate et al., 2022). Conversely, loss of Sod1 (or its chaperone Ccs1) should derepress *TDH3* regardless of the redox state, inhibiting the oxidative PPP. Corroborating this hypothesis, we also found the *TDH3* OEx strain to be a top-100 hit (**Suppl. Table S1)**

Lastly, disruption of the pyruvate dehydrogenase (PDH) complex, the entry point of pyruvate in the TCA cycle, was also found to be a KO hotspot. Two three-genes cliques and a number of additional genes were found at the top of the KO screen (**Fig. 4c**). These included *PDA1* and *LPD1*, which are parts of the PDH complex subunits E1 and E3, respectively **(Fig. 4d)**. Whatsmore, several genes encoding enzymes taking part in the synthesis of the cofactor lipoic acid (*HTD2, MCT1, LIP5)* and its loading (*AIM22, LIP2*) onto PDH subunit E2 were also found to be KO top hits.

In total, network analysis identified multiple gene cliques (**Suppl. Fig. S5**) and within six of these, the mechanistic interactions with CCM biosynthesis could be directly interpreted (**Fig. 4c**). Notably, we are the first to report the interaction between redox homeostasis and CCM synthesis, presumably through a common interaction with the oxidative PPP. Similarly, the network analysis suggested that improving riboflavin cofactor synthesis or introducing a pyruvate bottleneck by impairing the PDH complex should benefit CCM biosynthesis, corroborating previous observations (Blank et al., 2005; Pyne et al., 2018; Weber et al., 2017).

### Experimental validation of host:pathway interaction hits

In the previous section, CRI-SPA unveiled six gene cliques, which appear to interact with CCM synthesis. As a next step, we decided to validate hits sampled from these cliques (**Fig. 4d**) as well as a few hits from the cellular function category whose interaction with the pathway was less interpretable. We picked eight and six top hits in the OEx and KO screens, respectively. We also picked three low gene hits from the KO screen to further assess the correlation between CRI-SPA fluorescence score and CCM titers (**Suppl. Table S3)**.

For the KO hits, we purified strains from screening plates. The integrity of the pathway was verified by genotyping and the sequence of the unique barcodes identifying each gene deletion was verified by sequencing (Giaever et al., 2002). For the OEx hits, we placed each CDS under control of the strong *TEF2* promoter (Sun et al., 2012) and integrated the construct together with the CCM cassette into expression site XII-5 (Jessop-Fabre et al., 2016) of the parental BY4741 strain. To minimise the risk that the hit scores were the result of screening artefacts (e.g. plate, neighbour, agar effects, CCM diffusion in agar) we re-arrayed our validation strains on agar in 384 format interspersed with the control strain BY2. The fluorescence signal obtained in these conditions was well correlated with and less noisy than the original CRI-SPA score (PCC=0.70, p<10e-3, **Fig. 5a**). Again, we observed that the KO strains were generally less fluorescent than the OEx strains and that four of them (*LPD1, PDA1, AIM22, MET14*) were even less fluorescent than the WT. This simple rearraying could constitute a biosensor-mediated intermediate strain selection step, bridging high-throughput CRI-SPA (strains > 10,000s) to lower throughput HPLC (strains < 10s).

**Fig. 5.**
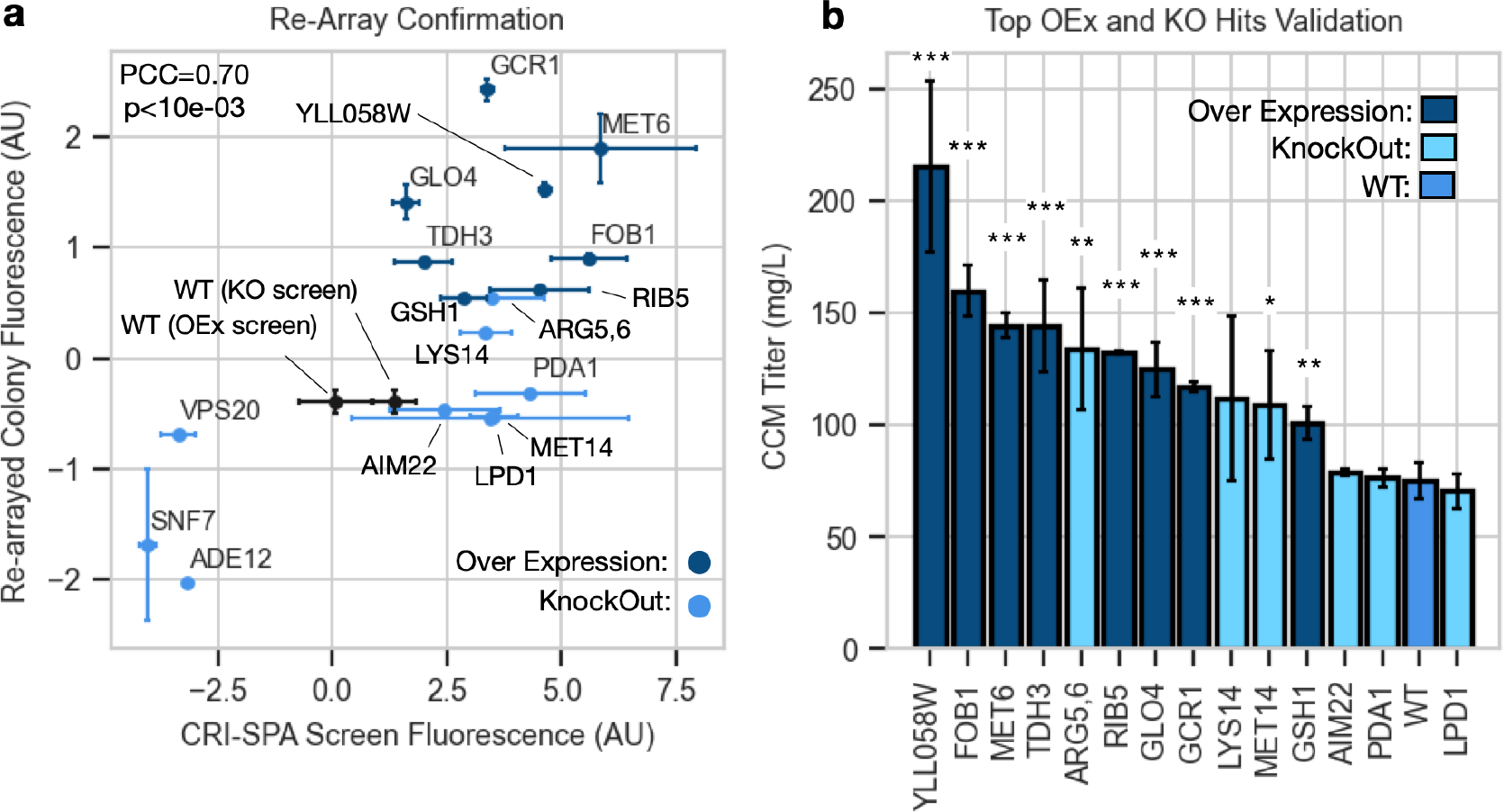
CRI-SPA hit validation. **a**) Scatter plot of fluorescences measured in the CRI-SPA screen and from plates of re-arrayed confirmation strains (PCC=0.70, p<10e-3). For the y-axis, the fluorescence of the re-arrayed plates were centred on the mean of their WT and scaled by the plate’s standard deviation. Points show the mean of four biological replicates and the error bars show their standard deviations. AU: arbitrary units. **b**) CCM titers for OEx top and bottom hits cultivated in liquid culture. Each bar represents the mean of 3-5 technical replicates. Biological replicate experiment for OEx hit is shown in **Suppl. Fig. S7**. Significant difference from WT is assessed with a two-sided t-test and indicated by: *<0.05, **<0.01, ***<0.001.

To verify that CRI-SPA fluorescence was representative of CCM accumulation, the bottom and top hits from the KO screen were cultivated in liquid Synthetic Complete (SC) medium for 72 h and CCM levels from individual supernatants were measured by HPLC. Here, we found a high correlation (PCC = >0.88) between the CRI-SPA fluorescence score and absolute HPLC measurements of CCM titers across multiple trials despite the batch-to-batch variation in the average CCM titers in between experiments **(Suppl. Fig. S6)**. This robust agreement between CRI-SPA score and CCM titers is a notable achievement given that CRI-SPA measures CCM levels with the help of a biosensor from colonies growing on solid medium.

To assess the ability of our screen to identify high CCM producers, we cultivated both OEx and KO top hits and measured their CCM levels. Importantly, we found that ten out of the fourteen tested top hits showed a significant increase in CCM titers compared to the BY2 reference strain (**Fig. 5b)**. Remarkably, all the selected OEx hits produced significantly more CCM than the reference strain. We also observed that OEx hits seemed to perform better than KO hits. This is in agreement with the observation that the OEx strains were more fluorescent than the KO strains (**Fig. 5a, Suppl. Fig. S3**) and that the difference between top hits and WT was larger in the OEx screen than in the KO screen (**Fig. 3**). The top four producers were all OEx hits with an improvement in CCM synthesis of 189-286%. The functions of the hits represented a variety of functional areas, from methionine (*YLL058W, MET6, MET14*), to arginine biosynthesis (*ARG5,6*) and the cryptic replication fork blocking gene (*FOB1*, **Suppl. Table S3**). Out of the four non-significant hits, three (*AIM22, LPD1* and *PDA1*) are involved in PDH function. This is a surprise given the number of top hits linked to PDH function found in the KO screen (**Fig. 4c**). We note however that these false positives could have been detected and removed from our analysis by the intermediate re-array step (**Fig. 5a**). Likewise, a biological replicate of the rearrayed OEx screen could not confirm the improved CCM titers of OEx top hits *RIB5* and *GSH1* (**Fig 5b, Suppl. Fig. S7**)

In summary, these validation results demonstrate the capacity of our CRI-SPA screen and data analysis to reliably identify new and diverse gene targets which positive interaction can be used in a metabolic engineering task, such as CCM biosynthesis in yeast cell factories.

## Discussion

The importance of host:pathway interactions has long been recognised (Cardinale and Arkin, 2012; de Lorenzo, 2011) and has motivated the development of a “host-aware” concept in synthetic biology (Boo et al., 2019). Focusing on the expression constructs themselves, successful strategies have sought to isolate them (Darlington et al., 2018; Lou et al., 2012; Meyer et al., 2015) or to minimise the burden they create on their host (Barajas et al., 2022; Ceroni et al., 2018). The importance of the host and its impact on the behaviour of engineered sequences is also increasingly being recognised (Calero and Nikel, 2019; Chan et al., 2023; Khan et al., 2020; Tas et al., 2021). For example, different host strains can show up to ten-fold differences in product titers (Strucko et al., 2015). Adapting the host to its construct opens a high-dimensional design space holding numerous exploitable phenotype maxima. This urges synthetic biologists to find the means to rapidly access and exploit these maxima.

ALE and genome-wide screens arguably search the host space for modifications improving a phenotype of interest. ALE screens the host genome for serendipitous edits and relies on fitness improvement to select the best ones (Mavrommati et al., 2022; Sandberg et al., 2019). Genome-wide screens introduce genetic perturbations, generally with CRISPR, recombineering or transposable elements (Cain et al., 2020; McCarty et al., 2020; Yilmaz et al., 2022), and screen and sort the edited library using massively parallelized reporter assays and next-generation sequencing to identify enriched edits (Bock et al., 2022; Cain et al., 2020). While both ALE and genome-wide screens have been extremely powerful at generating improved strains, the mechanistic learnings they generate are limited. For ALE, extracting the subset of beneficial mutations responsible for the trait improvement requires testing of individual and combinations of edits (Mundhada et al., 2017; Wang et al., 2020). Likewise, for genome-wide screens, a handful of best-performing strains usually dominate the top pool potentially obscuring other beneficial host:pathway interactions (Li et al., 2015; Savitskaya et al., 2019).

By editing each strain of an arrayed genome-wide library, CRI-SPA offers unprecedented functional tractability. The host space is systematically queried along every gene dimension for improvement of a desirable trait. The resulting dataset supports the building of hypotheses on the nature of the functional groups of hits identified. To facilitate this process, we propose in this study a computational strategy using STRING interaction networks, k-clique extraction, and GOEA annotation to identify functional gene clusters interacting with the synthesis of CCM, or any other analyte for which its quantification can be approximated by image analysis. This workflow has enabled the retrieval of a number of new gene clusters whose interactions could be deciphered as direct (e.g. riboflavin cofactor biosynthesis) and indirect (e.g. redox homeostasis effects on E4P precursor). We show that the top hits selected with the help of this *a priori* knowledge had a high true positive hit rate (10/14 >70%).

In its previous implementation, CRI-SPA relied on the yellowness of the plant pigment betaxanthin to measure strain productivity at high throughput (Cachera et al., 2023). This limited its application to a reduced number of analytes which are directly measurable via their colorimetric (e.g. beta-carotenes) or absorbance properties (e.g. *p*-coumaric acid), and which are frequently employed as optimisation targets in high-throughput screens (Zeng et al., 2020). In this work, the 19kb cassette encoded both the synthesis of CCM and a transcription factor-based biosensor (i.e. BenM MP17_D08) which, without further optimisation, allowed for the read-out of fluorescence directly from high-density agar arrays by image analysis. In the short-term future, we anticipate that the ease of biosensing, as implemented in this study, will open CRI-SPA’s applicability to a much wider range of small molecules and biologics which do not create a directly measurable phenotype (Dekker and Polizzi, 2017; Koch et al., 2019), and potentially the application of the CRI-SPA interaction screen as a general learning platform identifying debottlenecking strategies for genetically less tractable microorganisms.

## Materials and Methods

### Strains and Media

*E. coli* strains were cultivated in Luria-Bertani medium at 37°C containing 100 mg/L ampicillin to maintain plasmids. Yeast strains were propagated in YPD (10 g/L yeast extract, 20 g/L peptone, 20 g/L glucose) medium at 30°C. When appropriate, selection was performed with Nourseothricin (NTC) at 50 mg/L, Hygromycin at 100 mg/L and/or G418 at 100 mg/L in liquid medium and double these concentrations on solid medium.Ura3 counter selection was performed on solid SC medium supplemented with 1 g/L 5-fluoroorotic acid (5-FOA) and 30 mg/L uracil.

The YKO library was purchased from Transomic and the OEx TEF2pr-mCherry library was a kind gift from the Schuldiner lab (Weill et al., 2018). Strains and plasmids used in this study are available in **Suppl. Tables S4-S5**, respectively.

### General Cloning Procedure

All DNA sequences destined to cloning were amplified with Phusion U Hotstart (Thermofisher) using primers with USER compatible overhangs, assembled using USER cloning (Nour-Eldin et al., 2006) and transformed by heat shock in *E*.*coli* Dh5alpha. All plasmids inserts were sequenced and confirmed prior to yeast transformation. Yeast was transformed using the lithium acetate protocol (Gietz and Schiestl, 2007) and correct insertion was verified by PCR validation (DreamTaq Green, Thermo) both upstream and downstream of the integration site.

### Refactoring the CCM pathway

Repetitive promoters and terminators used to express the nine CDSs in the CCM pathway were replaced through three rounds of DNA assembly (**Suppl. Fig. S1**).The original promoter for BenM expression (REV1p) and the engineered pCYC_BenO driving yeGFP expression were left unmodified. In the first round, CDSs, promoters and terminators were fused into expression cassettes by USER assembly. This gave plasmids pCCM001-11. In the second round, multiple cassettes were fused and inserted into homology backbones containing flanking homology sequences. The homology sequences on the homology backbones were designed to insert the biosensor and the pathway together or on their own at site chromosomal site XII-5: pCCM021(XII-5Up:clone:HomoB), pCCM022 (HomoB:clone:XII-5Down), pCCM023 (HomoA:clone:HomoB) and pCCM024 (XII-5Up:clone:HomoA). Round two gave plasmids: pCCM012-16,18,20. In the third and final round, the joined expression cassettes were excised from pCCM012-16,18,20 and transformed and assembled into *S. cerevisiae* by homologous recombination. The USER primers used to PCR and fuse all CDSs, promoters, and terminators were automatically generated with our python script (https://github.com/pc2912/CRI-SPA_CCM).

### Gene knockout

The up and down homology sequences of the relevant ORF were fused to the kanMX cassette by USER cloning. USER primers were used to generate PCR fragments of the up- and downstream sequences of the ORFs using genomic DNA from BY4741 as template. The relevant PCR fragments were USER-fused to the kanMX cassette in *E. coli* and inserted into the USER cloning site of pCCM023, a modified version plasmid pCfB2909 lacking XII5-up and XII5-down homology sequences. The Up::kanMX::Dw gene-targeting substrates were released from purified plasmids by digestion with NotI prior to transformation into the relevant yeast strain which was selected on YPD containing G418.

### HPLC

Standards were prepared by diluting CCM (Sigma-Aldrich; 15992-5G-F) into deionized water to 200 mg/L and stirring until CCM was visibly dissolved. This dilution was heated at 95°C for 1 h and back diluted to obtain 0, 12.5, 25, 50, 100, and 200 mg/L dilutions which were used as standards for the calibration curve.

Strains were plated on selective SC agar medium (+NTC) to select for the CCM pathway. Individual colonies were picked in 2 mL of SC medium without antibiotic and grown overnight. The OD for each strain was adjusted to OD600 = 0.1 in 1 mL of the same medium in deep-well plates in experimental triplicates and grown for 72 h. After 72 h, the deep-well plate was spun at 3,000 *g* for 5 min. Next, 200 uL of supernatant was transferred to a 96-well plate which was tightly sealed and heated at 95°C for 1 h to kill any remaining cells. The plate was spun again at 3,000 *g* for 5 min and a ¼ to ⅛ dilution of the supernatant was used for HPLC quantification. Analysis was run on an Aminex HPX-87H ion exclusion column kept at 60 C using 1 mM H_2_SO_4_ as solvent injected at 0.6 ml/min. CCM was detected with a Thermo Fisher UV detector at 250 nm.

### Flow cytometry analysis

Single colonies were picked in 2 mL SC medium and grown overnight. Overnight cultures were diluted 50 times in 1 mL SC medium in 96-well plates and incubated in a shaking incubator set to 250 rpm, 30°C for 72 h. For CCM induction, SC medium was prepared containing CCM concentrations 710. 355, 177, 88, 8 and 0.8 mg/L. After cultivation, 1:4 water dilution of the cell cultures were analysed in a Quanteon flow cytometer (Novocyte). GFP fluorescence was measured using the B525 channel (emission: 488 nm, absorption 525/45 nm) for 12,000 events with size gate FSC-H > 3000. Flow cytometry data was manually gated and analysed with Python package Cytoflow (Teague, 2022).

### CRI-SPA pinning procedure

The pinning procedure was carried out with the Rotor (Singer Instruments) as previously described (Cachera et al., 2023). Briefly, the donor strain was grown overnight in liquid YPD with hygromycin to maintain the CRI-SPA vector to an OD between 0.6 and 2. Cells were washed twice with sterile deionised water to remove the antibiotic. A volume of 500 uL of the washed suspension was spread on YPD Singer Plus Plates which were left to dry in a laminar airflow cabinet until visibly dry. The library (in 384 format) was pinned in 4x4 quadruplicate (1,536 format) on top of the donor strain and incubated for 16 h at 30°C for mating. In our experiments we found that optimal mating efficiency was obtained when little donor library biomass was transferred to the mating plate. If the 384 library plates have big colonies, some colony biomass should be removed by a first uptake with 384 pads.

The mated arrays were replica pinned on the following agar substrates: YP + raffinose + hygromycin + antibiotic1 (24-48 h) > YP + galactose+ hygromycin + antibiotic1 (24 h) > SC 5-FOA (24-48 h) > SC + glucose + NTC + G418. Antibiotic1 is the library-specific antibiotic: G418 for YKO, and NTC for the OEx library. Glucose, raffinose and galactose were supplied at 20 g/L. All incubations were performed at 30°C. Images were acquired from the final SC + glucose + NTC + G418 plate after 48 h of incubation.

### Image acquisition and processing

For fluorescence measurements, screen plates were imaged with a Fusion-FX6 imager (Vilber) using the C480 light source, a F-535 Y2 filter and aperture option of 4 and an exposure time of 240 ms. Because lighting was not homogeneous within the imager, each plate was imaged with a rotation of 0, 90, 180, and 270 degrees. Colony fluorescence was then defined as the mean fluorescence extracted from the four pictures. For colony size, plates were imaged in white light with bottom lighting in a Phenobooth Imager (Singer instruments). Both raw colony size and fluorescence were extracted from plate pictures with the functions of the Pyphe Python package (Kamrad et al., 2020) as performed previously (Cachera et al., 2023).

### Data processing and normalisation

The experimental variance affecting colony fluorescence and size, was corrected using the same strategy developed for Synthetic Gene Array developed by the Boone and Andrews lab (Baryshnikova et al., 2010; Wagih et al., 2013). More specifically, the functions: Plate normalisation (N1), Spatial normalisation (N2), Row column effect (N3) and Jackknife filter (F4) were translated from R from (Wagih et al., 2013) and incorporated in our Python workflow. Raw colony fluorescence and colony size were processed in the following order: N1, N2, N3, F4, N1. After this processing, genes with less than three colony replicates were removed from our analysis. Scripts used for the analysis are available on our github repository (https://github.com/pc2912/CRI-SPA_CCM).

### Combining data from independent replicate screens

The raw data of two independent screens was passed through the following functions: N1, N2, N3, F4. At this step, N1 was applied so that the median of all plates from two screens were set to be equal. The data of the two screens was then pooled and genes with less than three colony replicates across the two screens were removed from our analysis.

### String database and networks

The STRING data was downloaded as file fi4932.protein.links.full.v11.5 from the database website (Szklarczyk et al., 2021). A group of genes was selected based on their ranking in the KO or OEx screens, for example the brightest 250 genes in the KO screen. An undirected graph was built representing genes as nodes and high confidence (confidence > 0.7) STRING associations as edges. Edges were coloured based on the type of association. For node colours, the data of the KO and OEx screens were merged in one dataframe to create dosage scores. If a gene readout was present only in one dataset, its score in the other dataset was assumed to be 0. A heuristic function was created to map the dosage score to an RGB value to maximise visible colour difference between genes on KO vs OEx plot (**Suppl. Fig. S4**).

Gene clusters were extracted by running NetworX’s k-clique communities function with k=3 (Palla et al., 2005). GOEA was run for the gene sets contained in each clusters with GOATOOLS (Consortium, 2006; Klopfenstein et al., 2018).

### KO Strains recovery and sequencing

For validation of the KO hits, strains were picked from the 1,536 screening plates and restreaked on NTC YPD plates. Fluorescence of the restreaks was verified and a representative colony for each hit was restreaked on a new NTC YPD plate. The integrity of the CCM-cassette was verified by amplifying its upstream and downstream integration sites. To validate the identity of the KO strain, the upstream and downstream barcode unique to each KO strain was amplified with primer pairs U1 + BTX.55 and U2+CCM.124 and Sanger sequenced. Both up and down barcodes were recovered for KO: *PDA1, ARG5,6, LYS14, MET14, LPD1, SNF7, ADE12* and *VPS20*. For *AIM22*, we only recovered the downstream barcode. Strains were cryostocked and used for further fluorescence and HPLC validations.

### OEx strains reverse engineering

The assembled CCM cassette was recovered by PCR from BY2 into two parts which were inserted into pCCM021 and pCCM039, resulting in plasmids pCCM042 and pCCM043, respectively. The ORFs of the selected OEx hits: *GCR1, MET6, GLO4, GSH1, TDH3, FOB1, RIB5* and *YLL058W* were amplified with their terminators from BY4741 genome and cloned downstream of TEF2p in pCCM40, resulting in plasmids pCCM055-59,61-63 (**Suppl. Table S5**). The insertion sites of pCCM021, pCCM039, and pCCM040 are flanked with matching homology sequences allowing for integration of their inserts into yeast genomic site XII-5 by homologous recombination. To create the OEx hits strain, the inserts of pCCM042 and pCCM043 and of either of pCCM055-59,61-63 were transformed together with pCF3050 in BY4741 harbouring Cas9 encoded on pHO38 and were selected on YPD + hygromycin + NTC + G418.

### Rearraying on 384 arrays

Because the OEx top hits were more fluorescent than the KO top hits, if arrayed on the same agar plate, the KO top hits’ fluorescence was “blinded” by that of the OEx top hits. To avoid this, we rearrayed the OEx hits and KO hits on two different agar arrays. The hit strains were pinned in square quadruplicates and interspersed with control strain BY2. BY2 was also placed at the edges of the array. In this way, all hit strains had BY2 as a direct neighbour and were not exposed to edge effects. Because of the high density of hits on the plates the “Boone and Andrews lab” correction (Wagih et al., 2013) was not applied. Rather, each plate data was standardised by subtracting the BY2 means and dividing by the plate’s standard deviation.

## Supporting information

Suppl. Table S2 Gene fluorescent ranking for the KO and OEx Screens

## Code and Data availability

The scripts used to design primers, extract the data from raw images, conduct analyses and reproduce the figures shown in this study are available on our Github repository: https://github.com/pc2912/CRI-SPA_CCM. The fluorescent and white light images for the OEx and one repeat of the KO screens are available to run the extraction and correction analyses. The raw and corrected fluorescence and colony size data for all screens mentioned in this study KO and OEx are also available on this repository.

## Acknowledgements

This study is supported by grants from the Novo Nordisk Foundation (NNF20CC0035580 and NNF19SA0035438) to MKJ and PP-YJC. We thank Prof. Lars Jul Jensen for his feedback on the progress of this work and his advice on using the STRING database. We thank Prof. Maya Schuldiner for kindly sending us the TEF2p SWAT overexpression library.

## Author contributions

P.C., U.H.M. and M.K.J. designed research. P.C., N.C.K and A.R. performed research. P.C. analyzed the data. T.S, U.H.M and M.K.J, supervised the research. P.C. and M.K.J. wrote the paper.

