## Supplementary Material for "Microbial cell factory optimisation using genome-wide host-pathway interaction screens"

**Supplementary figure S1 Cloning of the CCM cassette.**
**Supplementary figure S2: Biological replicate of experiments in figure 2.**
**Supplementary figure S3: Raw, uncorrected and corrected CRI-SPA data.**
**Supplementary figure S4: STRING Clusters for 250 most extreme gene groups.**
**Supplementary figure S5: K-clique clusters extracted from STRING networks and annotated with GOEA.**
**Supplementary Figure S6: CRI-SPA fluorescence score correlates with CCM titers in liquid culture.**
**Supplementary Figure S7: Biological replicate of experiment in figure 5.**

**Supplementary Table S1 Sourcing of parts used to build the CCM cassette.**
**Supplementary Table S2 Gene fluorescent ranking for the KO and OEx Screens**
**Supplementary Table S3 Gene hits selected for further testing.**
**Supplementary Table S4 Strains used in this study**
**Supplementary Table S5 Plasmids used in this study**
**Supplementary Table S6 Primers used in this study**

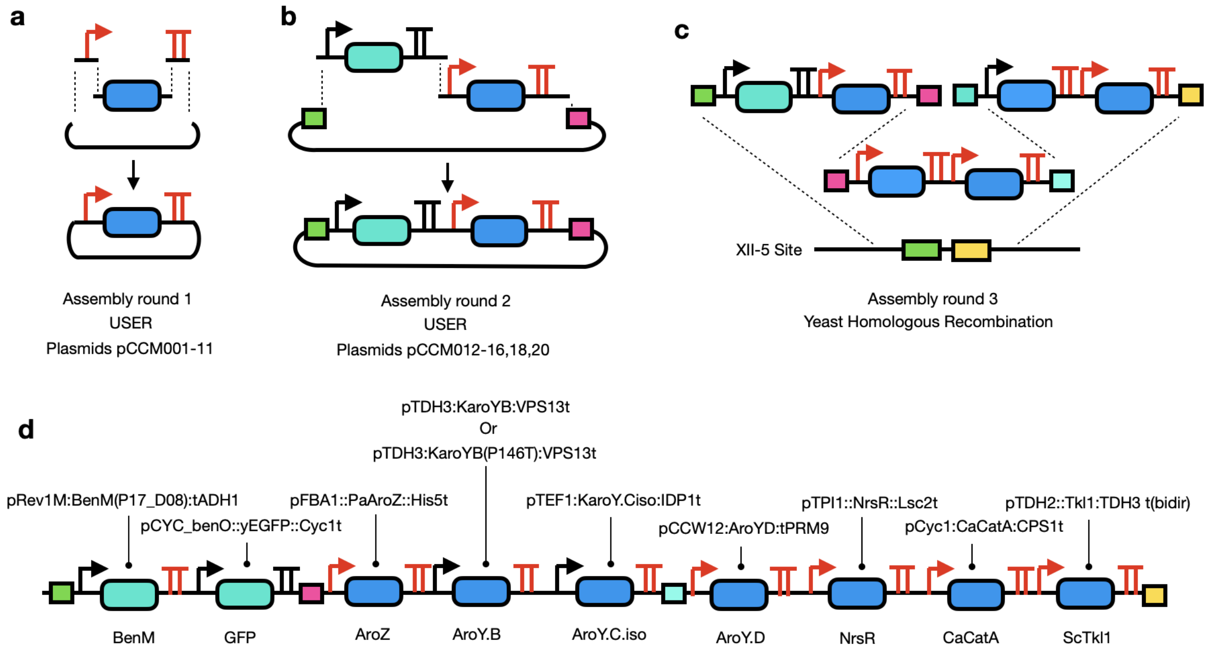


**Supplementary Figure S1. Refactoring the CCM cassette. a)** The CCM cassette in which repetitive promoters and terminators were replaced was built in three rounds of assembly. **a)** Individual parts (promoters, CDSs, terminators) were PCR amplified and USER-assembled into plasmids pCCM001-11. **b)** Expression cassettes from plasmids pCCM001-11 were PCR amplified and assembled into joined expression cassettes flanked by matching homology sequences for assembly in yeast. This yielded plasmids pCCM012-16,18, and 20. **c)** The joined expression cassettes from plasmids pCCM012-16,18,20 were excised and assembled by homologous recombination at integration site XII-5 in yeast. **d)** All parts in the final CCM expression cassette. The replaced promoters and terminators are marked in red. All parts and their sources are listed in **Supplementary Table S2**. All plasmids and primers used are listed in **Supplementary Tables S4-S5,** respectively.

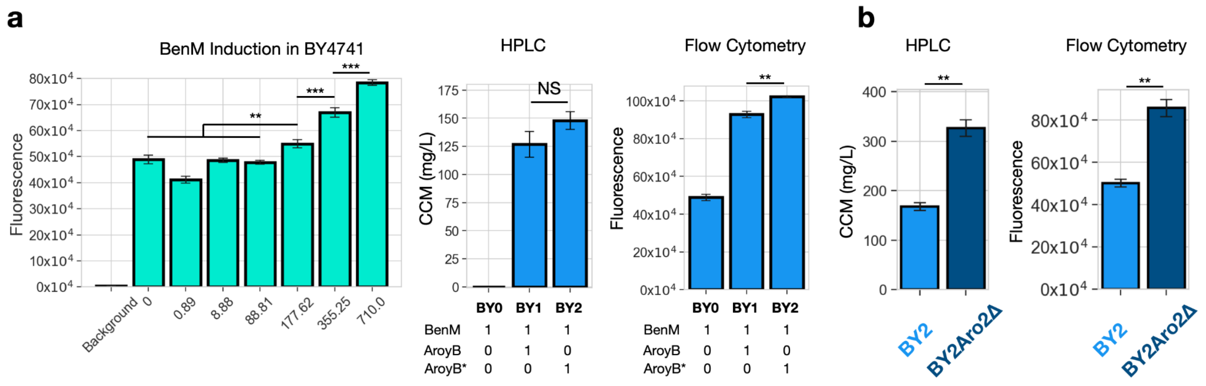


**Supplementary Figure S2: Biological replicate of experiments in Figure 2. a)** Left: induction of the biosensor in strain BY0. Centre and right: CCM titers and fluorescence measured by flow cytometry for BY0, BY1 and BY2. **b**) KO of positive control gene *Aro4* results in significant increase in CCM titers. The increase in CCM titers caused by *Aro2* causes a fluorescent change which can be detected by flow cytometry. For all graphs, significance was assessed with a two-sided t-test *<0.05, **<0.01, ***<0.001.

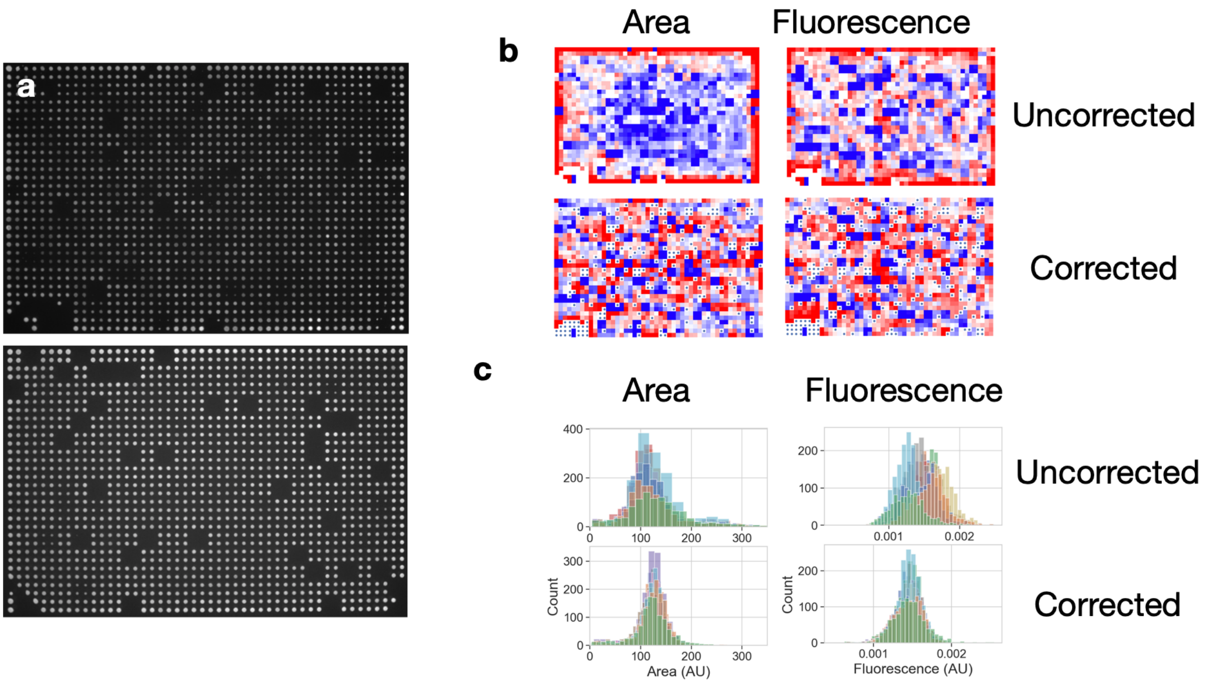

**Supplementary Figure S3. Raw, uncorrected and corrected CRI-SPA data. a)** Raw fluorescent images of screening plates produced by CRI-SPA for the YKO screen plate 2 (top) and the OEx screen (bottom). **b)** Uncorrected (top) and corrected (bottom) colony area and fluorescence data for YKO screen plate 2. Dots on the corrected plates mark empty spots and data points removed from our analysis. **c)** Distributions of the uncorrected (top) and corrected (bottom) data for each plate in the screen. Each colour is the distribution of data obtained for one plate.

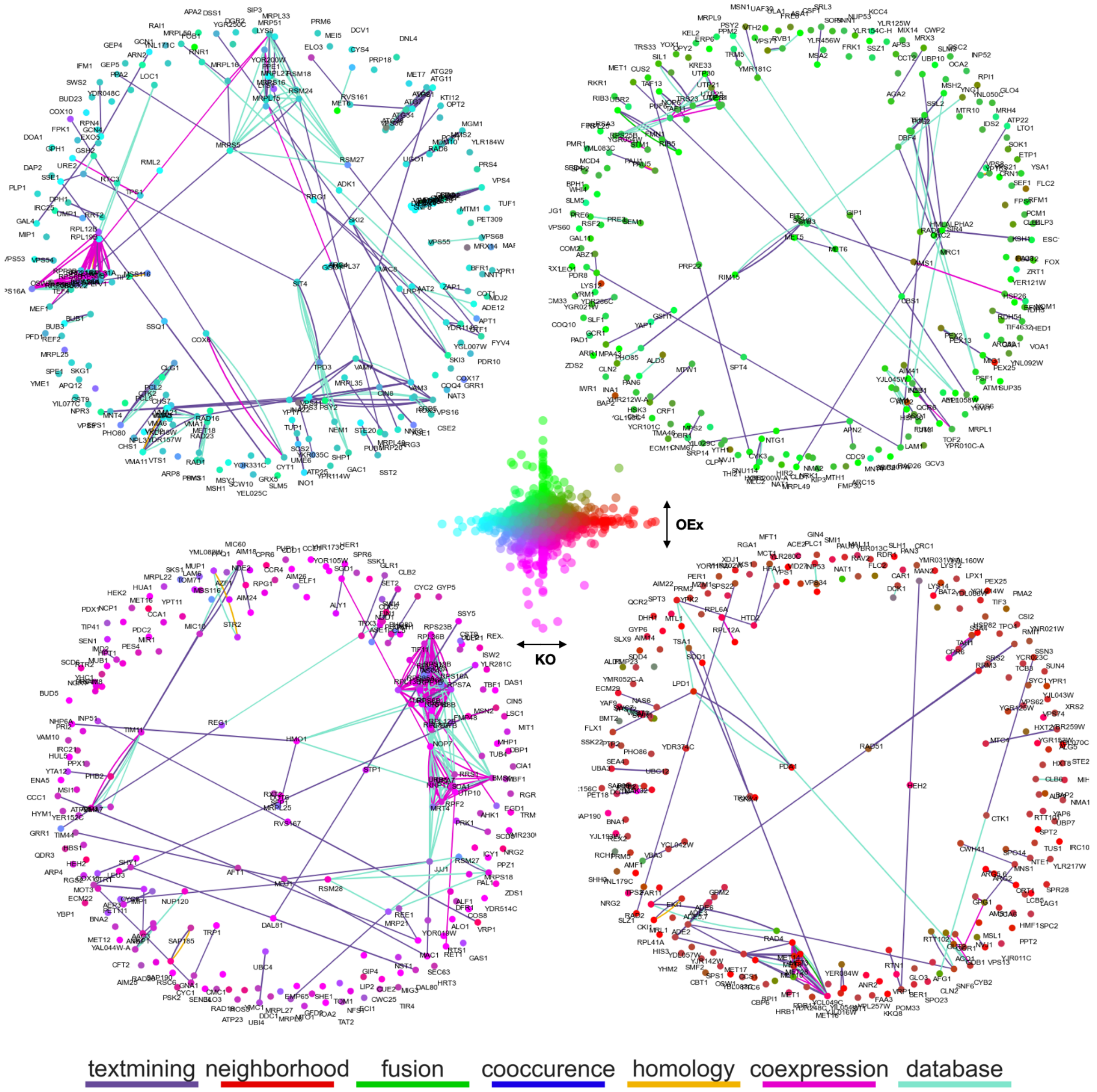


**Supplementary Figure S4. STRING Clusters for 250 most extreme gene groups.** Protein-protein interactions networks for the highest and lowest scoring genes in the YKO and the OEx screens. For the centre plot, each point is the mean fluorescence of a gene in the KO screen (x-axis) and the OEx screen (y-axis). Each gene was colour coded according to its x-y coordinate to map the effects of its KO and OEx on CCM synthesis in the rest of the figure. Graph nodes represent genes and are coloured according to their KO-OEx scores. For example, the top left graph represents the least fluorescent genes in the KO screen. Edges are STRING interactions of high confidence and coloured according to the bottom legend.

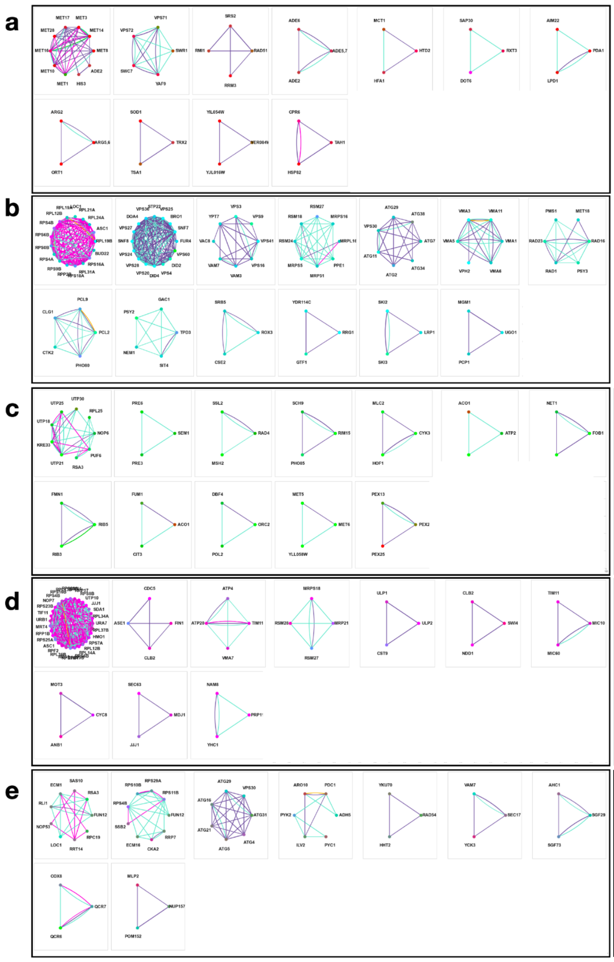

**a**

| | | | | | | |
|---|---|---|---|---|---|---|
| methionine biosynthetic process | chromatin remodeling | cellular response to DNA damage stimulus | purine nucleotide biosynthetic process | fatty acid Biosynthetic process | Rpd3L complex | Mitochond-rial pyruvate Dehydro-genase complex |
| arginine Biosynthetic process | antioxidant activity | None | protein folding | | | |

**b**

| | | | | | | |
|---|---|---|---|---|---|---|
| structural constituent of ribosome | endosome | fungal-type vacuole membrane | Mitochondrial small ribosomal subunit | autophagy | vacuolar acidification | DNA repair |
| cyclin-dependent protein kinase holoenzyme complex | protein phosphatase regulator activity | core mediator complex | None | RNA catabolic process | mitochondrial fusion | |

**c**

| | | | | | | |
|---|---|---|---|---|---|---|
| ribosome biogenesis | proteasome complex | DNA repair | protein kinase activity | cellular bud neck | Mitochon-drial nucleoid | nucleolus organization |
| riboflavin Biosynthetic process | Tricarbo-xylic acid cycle | mitotic DNA replication checkpoint signaling | None | peroxisomal membrane | | |

**d**

| | | | | | | |
|---|---|---|---|---|---|---|
| ribosome | spindle pole | proton trans-membrane transport | mitochondrial translation | SUMO-specific protease activity | None | Mitochond-rial crista junction |
| None | None | commitment complex | | | | |

**e**

| | | | | | | |
|---|---|---|---|---|---|---|
| ribosome biogenesis | Cytosolic small ribosomal subunit | phagophore assembly site | aromatic amino acid catabolic process to alcohol via Ehrlich pathway | None | fungal-type vacuole membrane | SAGA complex |
| respirasome | nuclear pore | | | | | |

**Supplementary Figure S5. K-clique clusters extracted from STRING networks and annotated with GOEA. a)** All K-clique gene clusters extracted from the gene networks shown in **Suppl. Fig. S4**. **a)**Top fluorescent group in KO screen, **b)** Bottom fluorescent group in KO screen. **c)** Top fluorescent group in OEx screen. **d)** Bottom fluorescent group in OEx screen. **e**) Cliques extracted for a STRING network made for 250 randomly selected genes. Left panels show the cliques, right panels show representative GO terms enriched in the corresponding clique.

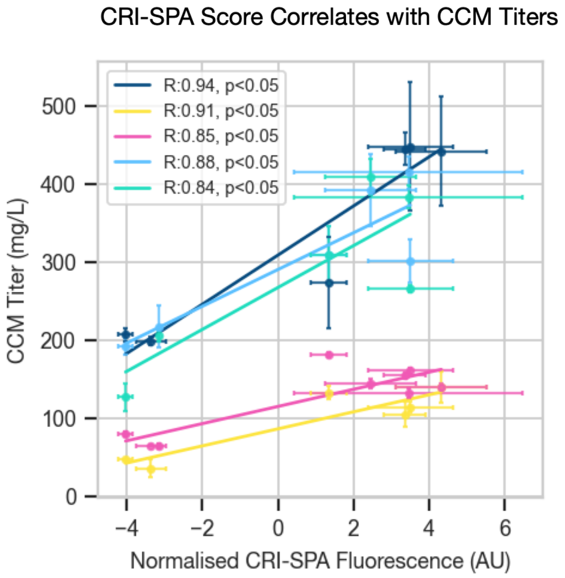


**Supplementary Figure S6. CRI-SPA fluorescence score correlates with CCM titers in liquid culture.** Scatter plot of fluorescences measured in the CRI-SPA screen and of CCM titers for top and bottom KO hits cultivated in SC medium for 72h. In the x-axis, each point is the mean fluorescence of three or more independent CRI-SPA colonies. In the y-axis, each point is the mean of three experimental replicates. Each colour represents a biological replicate performed on a different day. For both axes, the error bar is the standard deviation. R is the Pearson Correlation Coefficient between fluorescence intensities and CCM titers. AU = arbitrary units.

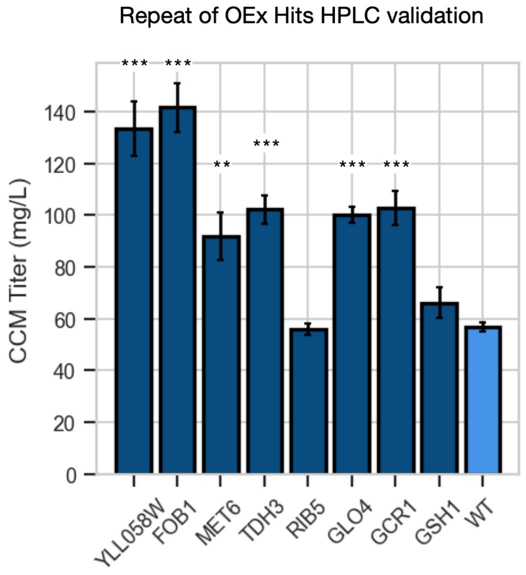


**Supplementary Figure S7. Biological replicates of experiment presented in Figure 5.** CCM titers for OEx top hits cultivated for 72h in liquid SC medium. Each bar represents the mean of three experimental replicates. Significant difference from WT is assessed with a two-sided t-test and indicated by: *<0.05, **<0.01, ***<0.001.

**Supplementary Table S1. Sourcing of parts used to build the CCM cassette.**

| **Part name** | **Primer Fw** | **Primer Rv** | **Description** | **Source/template** | **Reference** |
|---|---|---|---|---|---|
| pCYC_benO::yEGFP::tCyc1 | CCM001 | CCM002 | GFP expression cassette under BenM activated promoter | pMeLS0025 | Skjoedt et al |
| **pTDH2** | CMM005 | CCM006 | *S. cerevisiae* promoter | BY4741 genome | NA |
| **ScTkl1** | CMM007 | CCM008 | *S. cerevisiae TKL1* | pCFB1237 | Skjoedt et al |
| **tTDH3** | CCM009 | CCM010 | *S. cerevisiae* terminator | BY4741 genome | NA |
| **pFBA1** | CCM011 | CCM012 | *S. cerevisiae* promoter | BY4741 genome | NA |
| **PaAroZ** | CCM013 | CCM014 | *Podospora anserina's* dehydroshikimate dehydratase | pCFB1239 | Skjoedt et al |
| **tHis5** | CCM015 | CCM016 | *S. cerevisiae* terminator | BY4741 genome | NA |
| **pTPI1** | CCM017 | CCM018 | *S. cerevisiae* promoter | BY4741 genome | NA |
| **NTC_R** | CCM019 | CCM020 | Nourseothricin resistance | pCfB2193 | Stovicek et al. |
| **tLSC2** | CCM021 | CCM022 | *S. cerevisiae* terminator | BY4741 genome | NA |
| **pCYC1** | CMM023 | CMM024 | *S. cerevisiae* promoter | BY4741 genome | NA |
| **CaCatA** | CMM025 | CMM026 | *Candida albicans*' catechol 1,2-dioxygenase | pCFB1239 | Skjoedt et al |
| **tCPS1** | CMM027 | CMM028 | *S. cerevisiae* terminator | BY4741 genome | NA |
| **pTDH3:KpAroY.B** | CMM029 | CMM030 | *Klebsiella pneumonia* protocatechuic acid decarboxylase unit B under *S. cerevisiae's* pTdh3 promoter | pCfB1241 | Skjoedt et al |
| **pTDH3:KpAroY.B(P146T)** | CMM029 | CMM030 | Optimised *Klebsiella pneumonia* protocatechuic acid decarboxylase unit B under *S. cerevisiae's* pTdh3 promoter | Gblock | ED Jensen 2021 |
| **tVPS13** | CMM031 | CMM032 | *S. cerevisiae* terminator | BY4741 genome | NA |
| **pTef1:KpAroyC.iso** | CMM033 | CMM034 | *Klebsiella pneumonia* protocatechuic acid decarboxylase unit C under *S. Cerevisiae's* pTef1 promoter | pCFB1241 | Skjoedt et al |
| **tIDP1** | CMM035 | CMM036 | *S. cerevisiae* terminator | BY4741 genome | NA |
| **pCCW12** | CMM037 | CMM038 | *S. cerevisiae* promoter | BY4741 genome | NA |
| **KpAroY.D** | CCM039 | CCM040 | *Klebsiella pneumonia* protocatechuic acid decarboxylase unit D | pCfB1237 | Skjoedt et al |
| **tPRM9** | CCM041 | CCM042 | *S. cerevisiae* terminator | BY4741 genome | NA |

| **BenM MP17_D08** | CCM043 | CCM044 | Optimised version of BenM biosensor | pTS-117 | Snoek 2020 |
|---|---|---|---|---|---|

**Supplementary Table S2** Gene fluorescent ranking for the KO and OEx Screens

Find online

**Supplementary Table S3.** Gene hits selected for further testing.

| **Hit Name** | **Library** | **Normalised Fluorescence Score (Rank in Library)** | **Function** |
|---|---|---|---|
| **GCR1** | OEx | 3.39 (32th) | Transcriptional activator of genes involved in glycolysis |
| **MET6** | OEx | 5.85 (8th) | cobalamin-independent methionine synthase |
| **GLO4** | OEx | 1.62 (148th) | mitochondrial glyoxalase II, catalyzes the hydrolysis of s-d-lactoylglutathione into glutathione and d-lactate |
| **GSH1** | OEx | 2.85 (43th) | catalyzes the first step in glutathione (gsh) biosynthesis |
| **TDH3** | OEx | 2.00 (96th) | glyceraldehyde-3-phosphate dehydrogenase (GAPDH) |
| **FOB1** | OEx | 5.60 (10th) | nucleolar protein that binds the rDNA replication fork barrier site |
| **RIB5** | OEx | 4.53 (23th) | catalyzes the last step of the riboflavin biosynthesis |
| **YLL058W** | OEx | 4.64 (21th) | Inefficient homocysteine synthase |
| **LPD1** | KO | 3.45 (29th) | lipoamide dehydrogenase component (e3) of the pyruvate dehydrogenase |
| **LYS14** | KO | 3.35 (32nd) | transcriptional activator regulating lysine biosynthesis |
| **MET14** | KO | 3.53 (24rd) | adenylylsulfate kinase; required for sulfate assimilation and involved in methionine metabolism |
| **PDA1** | KO | 4.32 (11th) | e1 alpha subunit of the pyruvate dehydrogenase (pdh) complex |
| **AIM22** | KO | 2.44 (59th) | putative lipoate-protein ligase; required along with lip2 and lip5 for lipoylation of lat1p and kgd2p |
| **ARG5,6** | KO | 3.50 (256h) | catalyzes the 2nd and 3rd step in arginine biosynthesis |
| **VPS20** | KO | -3.36 (4586th) | part of the endosomal sorting complex required for transport of transmembrane |

| | | | |
|---|---|---|---|
| | | | proteins into the multivesicular body pathway to the lysosomal/vacuolar lumen |
| **ADE12** | KO | -3.15 (4577th) | catalyzes the first step during purine nucleotide biosynthesis |
| **SNF7** | KO | -4.03 (4592nd) | involved in the sorting of transmembrane proteins into the multivesicular body (mvb) pathway |

### Supplementary Table S4 Strains used in this study

| **Publication name** | **Parent** | **Genotype** | **Source** |
|---|---|---|---|
| **CRISPA-donor strain mat-alpha** | - | *MATα* CEN1-16::pGal1-*KlURA3can1-100 his3-11,15 leu2-3,112 LYS2 met17 trp1-1 ura3-1 RAD5* X-3::p*TEF1*-SpCas9-t*CYC1*-loxP-*KlLEU2 KlURA3(dw XII-5 IS)* | (Cachera et al., 2023) |
| **CRISPA-donor strain mat-a** | - | MATa CEN1-16::pGAL1-KlURA3can1-100 his3-11,15 leu2-3,112 LYS2 met17 trp1-1 ura3-1 RAD5 X-3::pTEF1-SpCas9-tCYC1-loxP-KlLEU2-loxP *KlURA3(dw XII-5 IS)* | (Cachera et al., 2023) |
| **BY4741** | - | MATa his3Δ1 leu2Δ0 met15Δ0 ura3Δ0 | (Baker Brachmann et al., 1998) |
| **CD1** | CRISPA-donor strain mat-alpha | *MATα* CEN1-16::pGal1-*KlURA3can1-100 his3-11,15 leu2-3,112 LYS2 met17 trp1-1 ura3-1 RAD5* X-3::p*TEF1*-SpCas9-t*CYC1*-loxP-*KlLEU2 KlURA3(dw XII-5 IS)* *XII-5::pCYC_BenO:yEGFP::BenM(MP17_D08)::PaAroZ::**AroY.B**::AroY.D:: NrsR: CaCatA::Tkl1* | This study |
| **CD2** | CRISPA-donor strain mat-alpha | *MATα* CEN1-16::pGal1-*KlURA3can1-100 his3-11,15 leu2-3,112 LYS2 met17 trp1-1 ura3-1 RAD5* X-3::p*TEF1*-SpCas9-t*CYC1*-loxP-*KlLEU2 KlURA3(dw XII-5 IS)* *XII-5::pCYC_BenO:yEGFP::BenM(MP17_D08)::PaAroZ::**AroY.B_P146T::AroY.D**:: NrsR:: CaCatA::Tkl1* | This study |
| **Sc_CCM052** | CRISPA-donor strain mat-a | *MATa CEN1-16::pGAL1-KlURA3can1-100 his3-11,15 leu2-3,112 LYS2 met17 trp1-1 ura3-1 RAD5 X-3::pTEF1-SpCas9-tCYC1-loxP-KlLEU2-loxP KlURA3(dw XII-5 IS)* *XII-5::pCYC_BenO:yEGFP::BenM(MP17_D08)::PaAroZ::**AroY.B**::AroY.D:: NrsR: CaCatA::Tkl1* | This study |
| **Sc_CCM053** | CRISPA-donor strain mat-a | *MATa CEN1-16::pGAL1-KlURA3can1-100 his3-11,15 leu2-3,112 LYS2 met17 trp1-1 ura3-1 RAD5 X-3::pTEF1-SpCas9-tCYC1-loxP-KlLEU2-loxP KlURA3(dw XII-5 IS)*<br><br>*XII-5::pCYC_BenO:yEGFP::BenM(MP17_D08)::PaAroZ::**AroY.B_P146T::AroY.Ciso::::AroY.D**:: NrsR:: CaCatA::Tkl1* | This study |
| **CD3** | Sc_CCM052 | *MATa CEN1-16::pGAL1-KlURA3can1-100 his3-11,15 leu2-3,112 LYS2 met17 trp1-1 ura3-1 RAD5 X-3::pTEF1-SpCas9-tCYC1-loxP-KlLEU2-loxP KlURA3(dw XII-5 IS)*<br><br>*XII-5::pCYC_BenO:yEGFP::BenM(MP17_D08)::PaAroZ::AroY.B**::AroY.Ciso:::AroY.D**:: **KanR**:: CaCatA::Tkl1* | This study |
| **CD4** | Sc_CCM053 | *MATa CEN1-16::pGAL1-KlURA3can1-100 his3-11,15 leu2-3,112 LYS2 met17 trp1-1 ura3-1 RAD5 X-3::pTEF1-SpCas9-tCYC1-loxP-KlLEU2-loxP KlURA3(dw XII-5 IS)* | This study |

| | | | |
|---|---|---|---|
| | | *XII-5::pCYC_BenO:yEGFP::BenM(MP17_D08)::PaAroZ::**AroY.B_P146T:**:AroY.Ciso:::AroY.D:: **KanR**: CaCatA::Tkl1* | |
| **BY0** | BY4741 | MATa his3Δ1 leu2Δ0 met15Δ0 ura3Δ0<br>*XII-5::pCYC_BenO:yEGFP::BenM(MP17_D08)* | This study |
| **BY1** | BY4741 | MATa his3Δ1 leu2Δ0 met15Δ0 ura3Δ0<br>*XII-5::pCYC_BenO:yEGFP::BenM(MP17_D08)::PaAroZ::**AroY.B**::AroY.Ciso::AroY.D::NrsR: CaCatA::Tkl1* | This study |
| **BY2** | BY4741 | MATa his3Δ1 leu2Δ0 met15Δ0 ura3Δ0<br><br>XII-5::pCYC_BenO:yEGFP::BenM(MP17_D08)::PaAroZ::**AroY.B_P146T::AroY.Ciso:::AroY.D**:: NrsR:: CaCatA::Tkl1 | This study |
| **BY2-Aro4Δ** | BY2 | MATa his3Δ1 leu2Δ0 met15Δ0 ura3Δ0 **Aro4Δ::KanMX**<br><br>XII-5::pCYC_BenO:yEGFP::BenM(MP17_D08)::PaAroZ::**AroY.B_P146T::AroY.Ciso::AroY.D**:: NrsR:: CaCatA::Tkl1 | This study |
| **BY2-Aro2Δ** | BY2 | MATa his3Δ1 leu2Δ0 met15Δ0 ura3Δ0 **Aro2Δ::KanMX**<br><br>XII-5::pCYC_BenO:yEGFP::BenM(MP17_D08)::PaAroZ::**AroY.B_P146T::AroY.Ciso:AroY.D**:: NrsR:: CaCatA::Tkl1 | This study |
| **OExHit_GCR1** | BY4741 (pHO38) | MATa his3Δ1 leu2Δ0 met15Δ0 ura3Δ0<br><br>**XII-5up**::pCYC_BenO:yEGFP::BenM(MP17_D08)::PaAroZ::**AroY.B_P146T**::AroY.Ciso:HomoB:AroY.D:KanR:: CaCatA::Tkl1::HomoC:Tef2p**GCR1** | This study |
| **OExHit_MET6** | BY4741 (pHO38) | MATa his3Δ1 leu2Δ0 met15Δ0 ura3Δ0<br><br>XII-5up::pCYC_BenO:yEGFP::BenM(MP17_D08)::PaAroZ::AroY.B_P146T::AroY.Ciso:HomoB:AroY.D:KanR:: CaCatA::Tkl1::HomoC:TEF2p:**MET6** | This study |
| **OExHit_:GLO4** | BY4741 (pHO38) | MATa his3Δ1 leu2Δ0 met15Δ0 ura3Δ0<br><br>XII-5up::pCYC_BenO:yEGFP::BenM(MP17_D08)::PaAroZ::AroY.B_P146T::AroY.Ciso:HomoB:AroY.D:KanR:: CaCatA::Tkl1::HomoC:TEF2p**:GLO4** | This study |
| **OExHit_GSH1** | BY4741 (pHO38) | MATa his3Δ1 leu2Δ0 met15Δ0 ura3Δ0<br><br>XII-5up::pCYC_BenO:yEGFP::BenM(MP17_D08)::PaAroZ::AroY.B_P146T::AroY.Ciso:HomoB:AroY.D:KanR:: CaCatA::Tkl1::HomoC:TEF2p:**GSH1** | This study |
| **OExHit_TDH3** | BY4741 (pHO38) | MATa his3Δ1 leu2Δ0 met15Δ0 ura3Δ0<br><br>XII-5up::pCYC_BenO:yEGFP::BenM(MP17_D08)::PaAroZ::AroY.B_P146T::AroY.Ciso:HomoB:AroY.D:KanR:: CaCatA::Tkl1::HomoC:TEF2p:**TDH3** | This study |
| **OExHit_FOB1** | BY4741 (pHO38) | MATa his3Δ1 leu2Δ0 met15Δ0 ura3Δ0<br><br>XII-5up::pCYC_BenO:yEGFP::BenM(MP17_D08)::PaAroZ::AroY.B_P146T::AroY.Ciso:HomoB:AroY.D:KanR:: CaCatA::Tkl1::HomoC:TEF2p:**FOB1** | This study |
| **OExHit_RIB5** | BY4741 (pHO38) | MATa his3Δ1 leu2Δ0 met15Δ0 ura3Δ0<br><br>XII-5up::pCYC_BenO:yEGFP::BenM(MP17_D08)::PaAroZ::AroY.B_P146T::AroY.Ciso:HomoB:AroY.D:KanR:: CaCatA::Tkl1::HomoC:TEF2p:**RIB5** | This study |

| | | | |
|---|---|---|---|
| **OExHit_YLL058W** | BY4741 (pHO38) | MATa his3Δ1 leu2Δ0 met15Δ0 ura3Δ0<br><br>XII-5up::pCYC_BenO:yEGFP::BenM(MP17_D08)::PaAroZ::AroY.B_P146T::AroY.Ciso:HomoB:AroY.D:KanR:: CaCatA::Tkl1::HomoC:TEF2p:**YLL058W** | This study |
| **KOhit_Pda1** | **Generated with CD2 x YKO CRI-SPA screen** | *MATα* CEN1-16::pGal1-*KlURA3can1-100 his3-11,15 leu2-3,112 LYS2 met17 trp1-1 ura3-1*<br>*RAD5* X-3::p*TEF1*-SpCas9-t*CYC1*-loxP-*KlLEU2 KlURA3(dw XII-5 IS)*<br>*XII-5::pCYC_BenO:yEGFP::BenM(MP17_D08)::PaAroZ::**AroY.B_P146T::AroY.D**:: NrsR:: CaCatA::Tkl1*<br>**Pda1Δ** | This study |
| **KOhit_ Arg5,6** | **Generated with CD2 x YKO CRI-SPA screen** | *MATα* CEN1-16::pGal1-*KlURA3can1-100 his3-11,15 leu2-3,112 LYS2 met17 trp1-1 ura3-1*<br>*RAD5* X-3::p*TEF1*-SpCas9-t*CYC1*-loxP-*KlLEU2 KlURA3(dw XII-5 IS)*<br>*XII-5::pCYC_BenO:yEGFP::BenM(MP17_D08)::PaAroZ::**AroY.B_P146T::AroY.D**:: NrsR:: CaCatA::Tkl1*<br>**Arg5,6Δ:KanMX** | This study |
| **KOhit_Lys14** | **Generated with CD2 x YKO CRI-SPA screen** | *MATα* CEN1-16::pGal1-*KlURA3can1-100 his3-11,15 leu2-3,112 LYS2 met17 trp1-1 ura3-1*<br>*RAD5* X-3::p*TEF1*-SpCas9-t*CYC1*-loxP-*KlLEU2 KlURA3(dw XII-5 IS)*<br>*XII-5::pCYC_BenO:yEGFP::BenM(MP17_D08)::PaAroZ::**AroY.B_P146T::AroY.D**:: NrsR:: CaCatA::Tkl1*<br>**Lys14Δ:KanMX** | This study |
| **KOhit_Met14** | **Generated with CD2 x YKO CRI-SPA screen** | *MATα* CEN1-16::pGal1-*KlURA3can1-100 his3-11,15 leu2-3,112 LYS2 met17 trp1-1 ura3-1*<br>*RAD5* X-3::p*TEF1*-SpCas9-t*CYC1*-loxP-*KlLEU2 KlURA3(dw XII-5 IS)*<br>*XII-5::pCYC_BenO:yEGFP::BenM(MP17_D08)::PaAroZ::**AroY.B_P146T::AroY.D**:: NrsR:: CaCatA::Tkl1*<br>**Met14Δ:KanMX** | This study |
| **KOhit_Lpd1** | **Generated with CD2 x YKO CRI-SPA screen** | *MATα* CEN1-16::pGal1-*KlURA3can1-100 his3-11,15 leu2-3,112 LYS2 met17 trp1-1 ura3-1*<br>*RAD5* X-3::p*TEF1*-SpCas9-t*CYC1*-loxP-*KlLEU2 KlURA3(dw XII-5 IS)*<br>*XII-5::pCYC_BenO:yEGFP::BenM(MP17_D08)::PaAroZ::**AroY.B_P146T::AroY.D**:: NrsR:: CaCatA::Tkl1*<br>**Lpd1Δ:KanMX** | This study |
| **KOhit_Snf7** | **Generated with CD2 x YKO CRI-SPA screen** | *MATα* CEN1-16::pGal1-*KlURA3can1-100 his3-11,15 leu2-3,112 LYS2 met17 trp1-1 ura3-1*<br>*RAD5* X-3::p*TEF1*-SpCas9-t*CYC1*-loxP-*KlLEU2 KlURA3(dw XII-5 IS)*<br>*XII-5::pCYC_BenO:yEGFP::BenM(MP17_D08)::PaAroZ::**AroY.B_P146T::AroY.D**:: NrsR:: CaCatA::Tkl1*<br>**Snf7Δ:KanMX** | This study |
| **KOhit_ Ade12** | **Generated with CD2 x YKO CRI-SPA screen** | *MATα* CEN1-16::pGal1-*KlURA3can1-100 his3-11,15 leu2-3,112 LYS2 met17 trp1-1 ura3-1*<br>*RAD5* X-3::p*TEF1*-SpCas9-t*CYC1*-loxP-*KlLEU2 KlURA3(dw XII-5 IS)*<br>*XII-5::pCYC_BenO:yEGFP::BenM(MP17_D08)::PaAroZ::**AroY.B_P146T::AroY.D**:: NrsR:: CaCatA::Tkl1*<br>**Ade12Δ:KanMX** | This study |
| **KOhit_Vps20** | **Generated with CD2 x YKO CRI-SPA screen** | *MATα* CEN1-16::pGal1-*KlURA3can1-100 his3-11,15 leu2-3,112 LYS2 met17 trp1-1 ura3-1*<br>*RAD5* X-3::p*TEF1*-SpCas9-t*CYC1*-loxP-*KlLEU2 KlURA3(dw XII-5 IS)*<br>*XII-5::pCYC_BenO:yEGFP::BenM(MP17_D08)::PaAroZ::**AroY.B_P146T::AroY.D**:: NrsR:: CaCatA::Tkl1*<br>**Vps20Δ:KanMX** | This study |

| | | | |
|---|---|---|---|
| **KOhit_Aim22** | **Generated with CD2 x YKO CRI-SPA screen** | *MATα* CEN$_{1\text{-}16}$::pGal1-*KlURA3can1-100 his3-11,15 leu2-3,112 LYS2 met17 trp1-1 ura3-1* *RAD5* X-3::p*TEF1*-SpCas9-t*CYC1*-loxP-*KlLEU2 KlURA3(dw XII-5 IS)* *XII-5::pCYC_BenO:yEGFP::BenM(MP17_D08)::PaAroZ::**AroY.B_P146T::AroY.D**:: NrsR:: CaCatA::Tkl1* **Aim22Δ:KanMX** | This study |

### Supplementary Table S5 Plasmids used in this study

| Name | Backbone | Description | Type |
|---|---|---|---|
| pCCM001 | pCFB2909 | pCYC_benO::yEGFP::Cyc1t | CCM Cassette Assembly Round 1 |
| pCCM003 | pCFB2909 | pTDH2::Tkl1:TDH3 t(bidir) | CCM Cassette Assembly Round 1 |
| pCCM004 | pCFB2909 | pFBA1::PaAroZ::His5t | CCM Cassette Assembly Round 1 |
| pCCM005 | pCFB2909 | pTPI1::NrsR::Lsc2t | CCM Cassette Assembly Round 1 |
| pCCM006 | pCFB2909 | pCyc1:CaCatA:CPS1t | CCM Cassette Assembly Round 1 |
| pCCM007 | pCFB2909 | pTDH3:KaroYB:VPS13t | CCM Cassette Assembly Round 1 |
| pCCM008 | pCFB2909 | pTEF1:KaroY.Ciso:IDP1t | CCM Cassette Assembly Round 1 |
| pCCM009 | pCFB2909 | pCCW12:AroYD:tPRM9 | CCM Cassette Assembly Round 1 |
| pCCM010 | pCFB2909 | tADH1:BenM(P17_D08):pRev1M | CCM Cassette Assembly Round 1 |
| pCCM011 | pCFB2909 | pTDH3:KaroYB(P146T):VPS13t | CCM Cassette Assembly Round 1 |
| pCCM012 | pCCM021 | XII-5up::PaAroZ::AroY.B:: AroY.Ciso:HomoB | CCM Cassette Assembly Round 2 for pathway alone |

| | | | |
|---|---|---|---|
| pCCM013 | pCCM021 | XII-5up::PaAroZ::AroY.B_P146T:: AroY.Ciso:HomoB | CCM Cassette Assembly Round 2 for pathway alone |
| pCCM014 | pCCM023 | HomoA::PaAroZ::AroY.B:: AroY.Ciso:HomoB | CCM Cassette Assembly Round 2 for pathway + sensor |
| pCCM015 | pCCM023 | HomoA::PaAroZ::AroY.B_P146T:: AroY.Ciso:HomoB | CCM Cassette Assembly Round 2 for pathway + sensor |
| pCCM016 | pCCM022 | HomoB: AroY.D:: NrsR:CaCatA::Tkl1::XII-5dwn | CCM Cassette Assembly Round 2 for pathway alone and for pathway + sensor |
| pCCM018 | pCCM025 | XII-5up::yEGFP::BenM(MP17_D08):XII-5dwn | CCM Cassette Assembly Round 2 Sensor alone |
| pCCM020 | pCCM024 | XII-5up::yEGFP::BenM(MP17_D08):HomoA | CCM Cassette Assembly Round 2 for pathway + sensor |
| pCCM021 | pCFB2909 | XII-5Up:clone:HomoB | Homology Backbone |
| pCCM022 | pCFB2909 | HomoB:clone:XII-5Down | Homology Backbone |
| pCCM023 | pCFB2909 | HomoA:clone:HomoB | Homology Backbone |
| pCCM024 | pCFB2909 | XII-5Up:clone:HomoA | Homology Backbone |
| pCCM026 | pCCM023 | Aro2:KanMX KO | Used to KO Aro2 |
| pCCM027 | pCCM023 | Aro4:KanMX KO | Used to KO Aro4 |
| pCCM039 | pCCM022 | HomoB:clone:HomoC | Used for OEx strains cloning |
| pCCM040 | pCCM022 | HomoC::clone::XII5 | Used for OEx strains cloning |
| pCCM042 | pCCM021 | XII-5up::pCYC_BenO:yEGFP::BenM(MP17_D08)::PaAroZ::AroY.B_P146T::AroY.Ciso:HomoB | Used for OEx strains cloning |
| pCCM043 | pCCM039 | HomoB: AroY.D:KanR:: CaCatA::Tkl1::HomoC | Used for OEx strains cloning |

| | | | |
|---|---|---|---|
| pCCM044 | pCCM021 | Tp1_P:KanR:LSC2_T | Used to switch the NTC resistance on to Kan resistance in CD strains |
| pCCM055 | pCCM040 | HomoC:: pTEF2::GCR1 ::XII-5Dw | Used for OEx strains cloning |
| pCCM056 | pCCM040 | HomoC:: pTEF2::MET6 ::XII-5Dw | Used for OEx strains cloning |
| pCCM057 | pCCM040 | HomoC:: pTEF2::GLO4 ::XII-5Dw | Used for OEx strains cloning |
| pCCM058 | pCCM040 | HomoC:: pTEF2::GSH1 ::XII-5Dw | Used for OEx strains cloning |
| pCCM059 | pCCM040 | HomoC:: pTEF2::TDH3 ::XII-5Dw | Used for OEx strains cloning |
| pCCM061 | pCCM040 | HomoC:: pTEF2::FOB1 ::XII-5Dw | Used for OEx strains cloning |
| pCCM062 | pCCM040 | HomoC:: pTEF2::RIB5 ::XII-5Dw | Used for OEx strains cloning |
| pCCM063 | pCCM040 | HomoC:: pTEF2::YLL058W ::XII-5Dw | Used for OEx strains cloning |
| pHO029 | | pSNR52-gRNA(XII-5)-tSUP4 | CRI-SPA vector with |
| pHO38 | | CRISPR-Cas9 (Hyg) | |
| pCFB3050 | | pSNR52-gRNA(XII-5)-tSUP4 | |

### Supplementary Table S6 Primers used in this study

| **Primer Name** | **Bases** | **Tm (c)** | **User overhang** | **Template** |
|---|---|---|---|---|
| **CCM.001** | CGTGCGAUGATCCAGGCAACTTTAGTGC | 52.81 | GV1 | pMeLS0025(pCYC_benO::yEGFP::Cyc1t) |
| **CCM.002** | CACGCGAUTTCAGCTGACGCGATCT | 50.11 | GV2 | pMeLS0025(pCYC_benO::yEGFP::Cyc1t) |
| **CCM.003** | CACGCGAUGAATGCGGCCGCTT | 50.01 | GV2 | pMeLS0076(Rev1p::BenM(H110R, F211V, Y286N)) |
| **CCM.004** | CGTGCGAUAGCGGATAACAATTTCACACA | 51.41 | GV1 | pMeLS0076(Rev1p::BenM(H110R, F211V, Y286N)) |

| | | | | |
|---|---|---|---|---|
| **CCM. 005** | AAGAGGGCUTTTGTTTTGTTTGTTTGTGTGATGAATTTAA | 53. 92 | U1 | pTDH2 |
| **CCM. 006** | CGTGCGAUATCTAGATCAGAGGGTGGT | 50. 32 | GV1 | pTDH2 |
| **CCM. 007** | AGCCTGTGUTCAGAAAGCTTTTTTCAAAGGA | 49. 14 | U2 | pCFB1237_ScTkl1 |
| **CCM. 008** | AGCCCTCTUACAATGACTCAATTCACTGACATTGAT | 53. 72 | U1 | pCFB1237_ScTkl1 |
| **CCM. 009** | CACGCGAUTCTGCAGGTAGGGAAAGA | 51. 07 | GV2 | ANTE113_Tdh3bidir |
| **CCM. 010** | ACACAGGCUGTGAATTTACTTTAAATCTTGCATTTAA | 50. 04 | U2 | ANTE113_Tdh3bidir |
| **CCM. 011** | AAGAGGGCuTTTGAATATGTATTACTTGGTTATG | 48. 18 | U1 | FBA1p |
| **CCM. 012** | CGTGCGAuACTGGTAGAGAGCGACTTT | 50. 78 | GV1 | FBA1p |
| **CCM. 013** | AGCCTGTGuTCACAAAGCAGCTGACAAAG | 51. 02 | U2 | pCFB1239_PaAroZ |
| **CCM. 014** | AGCCCTCTuACAATGCCATCCAAGTTGG | 52. 12 | U1 | pCFB1239_PaAroZ |
| **CCM. 015** | ACACAGGCuATAGATTAATTTAAACAGTATATGTACAGTTT | 51. 49 | U2 | His5t |
| **CCM. 016** | CACGCGAuGTAACAATATCATGAGACCTTTTATA | 49. 64 | GV2 | His5t |
| **CCM. 017** | AAGAGGGCuTTTTAGTTTATGTATGTGTTTTTTGTAGT | 51. 25 | U1 | TPI1p |
| **CCM. 018** | CGTGCGAuAAGGATGAGCCAAGAATAAGG | 51. 72 | GV1 | TPI1p |
| **CCM. 019** | AGCCCTCTuACCATGGGTACCACTCT | 51. 25 | U1 | pCfB2193_NatMX |
| **CCM. 020** | AGCCTGTGuTTAGGGGCAGGGCA | 51. 63 | U2 | pCfB2193_NatMX |
| **CCM. 021** | CACGCGAuAAAATTAAAAAAAAAAAAAAGAAATTTTTTCC | 48. 92 | GV2 | LSC2t |
| **CCM. 022** | ACACAGGCuGCTTCTCGAGAAAAACAAAAGAGTTA | 53. 07 | U2 | LSC2t |
| **CCM. 023** | CGTGCGAuCAGCATTTTCAAAGGTGT | 45. 87 | GV1 | pCYC1 |
| **CCM. 024** | AAGAGGGCuTATTAATTTAGTGTGTGTATTTGTGTT | 50. 24 | U1 | pCYC1 |
| **CCM. 025** | AGCCCTCTuACAATGTCCCAAGCTTTCA | 50. 21 | U1 | pCFB1239CaCatA |

| | | | | |
|---|---|---|---|---|
| **CCM. 026** | AGCCTGTGuTTACAACTTGATTTCAGCATCTTG | 50. 74 | U2 | pCFB1239CaCatA |
| **CCM. 027** | ACACAGGCuGCGCAATGATTGAATAGTCAAA | 50. 74 | U2 | CPS1t |
| **CCM. 028** | CACGCGAuGATTTGACACTTGATTTGACACTTCTTT | 53. 48 | GV2 | CPS1t |
| **CCM. 029** | AGCCTGTGuTCATTCAATTTCTTGAGCGAATTG | 50. 87 | U2 | pCfB1241_TDH3p:KpAroY .B |
| **CCM. 030** | CGTGCGAuGCTATAAAAAACACGCTTTTTCA | 50. 16 | GV1 | pCfB1241_TDH3p:KpAroY .B |
| **CCM. 031** | CACGCGAuGCGCGCTGCGGA | 51. 74 | GV2 | VPS13t |
| **CCM. 032** | ACACAGGCuTCACATATGAAAGTATATACCCG | 50. 63 | U2 | VPS13t |
| **CCM. 033** | CGTGCGAuGCACACACCATAGCTTCAA | 51. 31 | GV1 | pCFB1241_Tef1p_KpAroy C.iso |
| **CCM. 034** | AGCCTGTGuTTACTTAGCGGAACCTTGATT | 51. 33 | U2 | pCFB1241_Tef1p_KpAroy C.iso |
| **CCM. 035** | ACACAGGCuTCGAATTTACGTAGCCCAATCTA | 54. 23 | U2 | IDP1t |
| **CCM. 036** | CACGCGAuGATGGTAATGATCCGAACTTGG | 53. 28 | GV2 | IDP1t |
| **CCM. 037** | AAGAGGGCuTATTGATATAGTGTTTAAGCGAATGA | 50. 87 | U1 | CCW12p |
| **CCM. 038** | CGTGCGAuAAAGAAACTTAATACGTTATGCC | 49. 65 | GV1 | CCW12p |
| **CCM. 039** | AGCCCTCTuACAATGATCTGTCCAAGATGC | 50. 99 | U1 | pCfB1237_KpAroY.D |
| **CCM. 040** | AGCCTGTGuTTATCTCTTATCTTCTGGCAATAATG | 51. 11 | U2 | pCfB1237_KpAroY.D |
| **CCM. 041** | ACACAGGCuACAGAAGACGGGAGACAC | 51. 44 | U2 | PRM9t |
| **CCM. 042** | CACGCGAuATTTTCAACATCGTATTTTCCGA | 50. 34 | GV2 | PRM9t |
| **CCM. 043** | CGTGCGAuGAATGCGGCCGCTT | 50. 01 | GV1 | pMeLS0076(Rev1p::BenM (H110R, F211V, Y286N)) |
| **CCM. 044** | CACGCGAuAGCGGATAACAATTTCACACA | 51. 41 | GV2 | pMeLS0076(Rev1p::BenM (H110R, F211V, Y286N)) |
| **CCM. 045** | AAGAGGGCuctcgctgaggacttaatgc | | U1 | pCFB2909 |

| | | | | |
|---|---|---|---|---|
| **CCM. 046** | AGCCCTCTugaatgcgtgcgatcgcgtgcattcACTGAATACAGGCTAGTGAATGCTCTCGGTCTTGTACGTGTGCGATTGACagattaagacctcagcgc | | U1 | pCFB2909 |
| **CCM. 047** | AGCCCTCTugaatgcgtgcgatcgcgtgcattcagtgctgaggcattaatgc | | U1 | pCFB2909 |
| **CCM. 048** | AAGAGGGCuGTCAATCGCACACGTACAAGACCGAGAGCATTCACTAGCCTGTATTCAGTtgattaaaccctcagcgcg | | U1 | pCFB2909 |
| **CCM. 049** | AAGAGGGCuGTTGTCGTGCCACCGTTACCTTCGTGAGTATTCAATCTCATAGCCGAGTAtgattaaaccctcagcgcg | | U1 | pCFB2909 |
| **CCM. 050** | AGCCCTCTugaatgcgtgcgatcgcgtgcattcTACTCGGCTATGAGATTGAATACTCACGAAGGTAACGGTGGCACGACAACcagattaagacctcagcgc | 102 | U1 | pCFB2909 |
| **CCM. 012** | CGTGCGAuACTGGTAGAGAGCGACTTT | 50.78 | GV1 | pCCM004(pFBA1:PaAroZ:tHis5) |
| **CCM. 051** | AAGAGGGCuGTAACAATATCATGAGACCTTTTATAG | 50.67 | U1 | pCCM004(pFBA1:PaAroZ:tHis5) |
| **CCM. 052** | AGCCCTCTuATAAAAAACACGCTTTTTCAGTTC | 50.2 | U1 | pCCM007(pTDH3:KpAroyB:tVPS13) |
| **CCM. 053** | AGCCTGTGuGCGCGCTGCGGA | 51.74 | U2 | pCCM007(pTDH3:KpAroyB:tVPS13) |
| **CCM. 054** | ACACAGGCuGCACACACCATAGCTTCA | 50.54 | U2 | pCCM008(pTEF1:Aroy.Ciso:tIDP1) |
| **CCM. 036** | CACGCGAuGATGGTAATGATCCGAACTTGG | | GV2 | pCCM008(pTEF1:Aroy.Ciso:tIDP1) |
| **CCM. 038** | CGTGCGAuAAAGAAACTTAATACGTTATGCC | | GV1 | pCCM009(pCCW12:AroYD:tPRM9) |
| **CCM. 055** | AAGAGGGCuATTTTCAACATCGTATTTTCCGA | 50.34 | U1 | pCCM009(pCCW12:AroYD:tPRM9) |
| **CCM. 056** | AGCCCTCTuTAAGGATGAGCCAAGAATAAGG | 52.15 | U1 | pCCM005(pTPI1:NrsR:LSC2t) |
| **CCM. 057** | AGCCTGTGuAAAATTAAAAAAAAAAAAAAGAAATTTTTTCCA | 50.07 | U2 | pCCM005(pTPI1:NrsR:LSC2t) |
| **CCM. 058** | ACACAGGCuTCAGCATTTTCAAAGGTGTGT | 51.15 | U2 | pCCM006(pCyc1:CaCatA:CPS1t) |
| **CCM. 059** | AGGTGGCAuGATTTGACACTTGATTTGACACTT | 50.4 | U3 | pCCM006(pCyc1:CaCatA:CPS1t) |

| | | | | |
|---|---|---|---|---|
| **CCM. 060** | ATGCCACCuATCTAGATCAGAGGGTGGT | 50.32 | U3 | pCCM003(pTDH2:Tkl1:TDH3t) |
| **CCM. 009** | CACGCGAuTCTGCAGGTAGGGAAAGA | 51.07 | GV2 | pCCM003(pTDH2:Tkl1:TDH3t) |
| **CCM. 001** | CGTGCGAuGATCCAGGCAACTTTAGTGC | 52.81 | GV1 | pCCM001(pCYC_benO:yEGFP:Cyc1t) |
| **CCM. 061** | AAGAGGGCuTTCAGCTGACGCGATCT | 50.11 | U1 | pCCM001(pCYC_benO:yEGFP:Cyc1t) |
| **CCM. 062** | CACGCGAuGCGACCTCATGCTATACC | 51.71 | GV2 | pCCM002(tADH1::BenM::pRev1) and pCCM010(tADH1::BenM(MP17_D08)::pRev1 |
| **CCM. 063** | AGCCCTCTuAGCGGATAACAATTTCACACA | 51.41 | U1 | pCCM002(tADH1::BenM::pRev1) |
| **CCM. 063B** | AGCCCTCTuTTCTTAGGCACAACAATATTTATAAA | | U1 | pCCM010(tADH1::BenM(MP17_D08)::pRev1 |
| **CCM. 064** | CGTGCGAuTTACCAATTTGGTGGTTCAG | 50.21 | GV1 | pMeLS0076(Rev1p::BenM_MP17_D08) |
| **CCM. 065** | CACGCGAuTTCTTAGGCACAACAATATTTATAAA | 50.1 | GV2 | pMeLS0076(Rev1p::BenM_MP17_D08) |
