## Supplementary material for "Microbial cell factory optimisation using genome-wide host-pathway interaction screens": Figures & Suppl. Figures

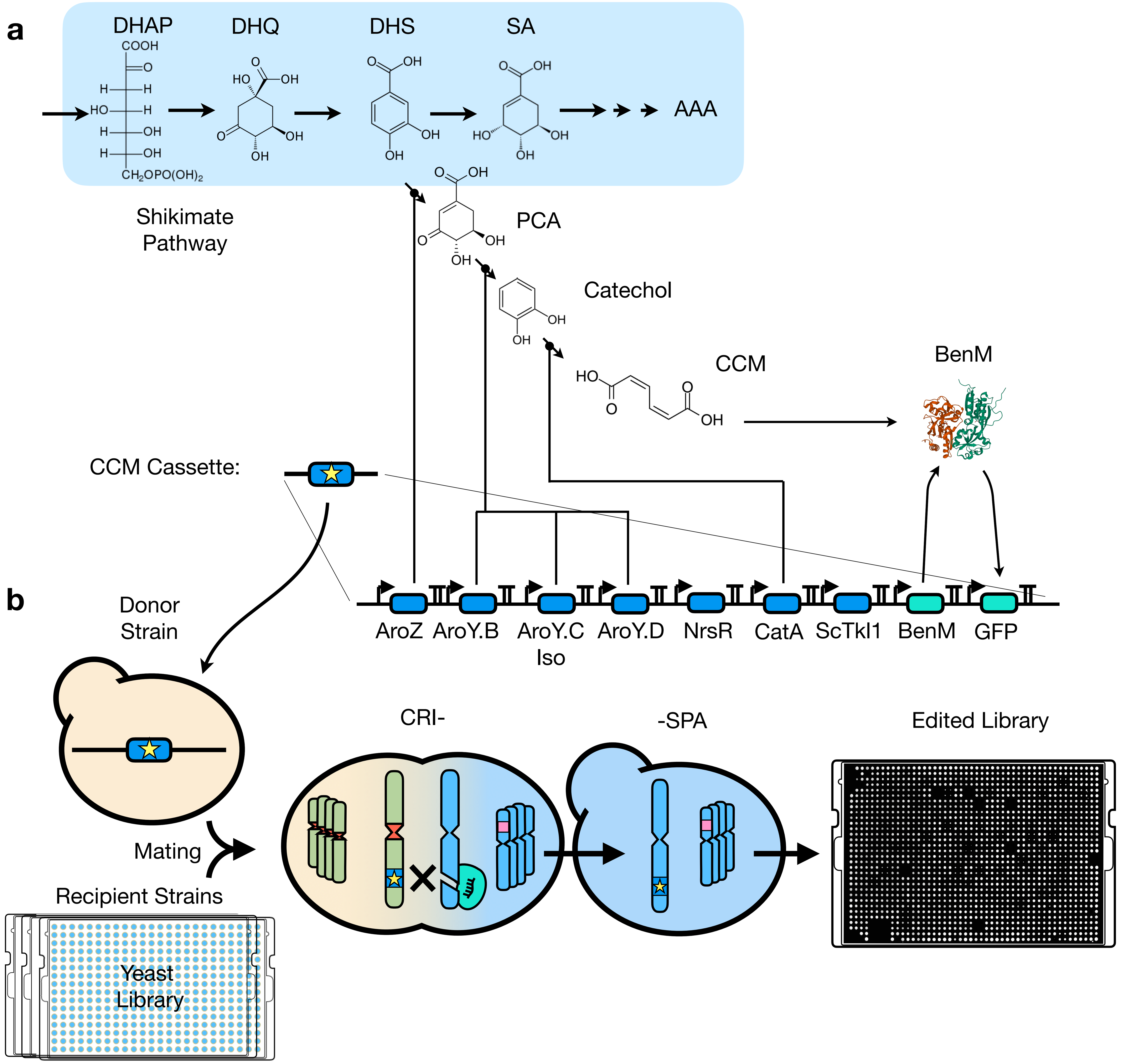


**Fig. 1. Overview of the design of the CRI-SPA screen for identification of host genes interacting with CCM synthesis.**

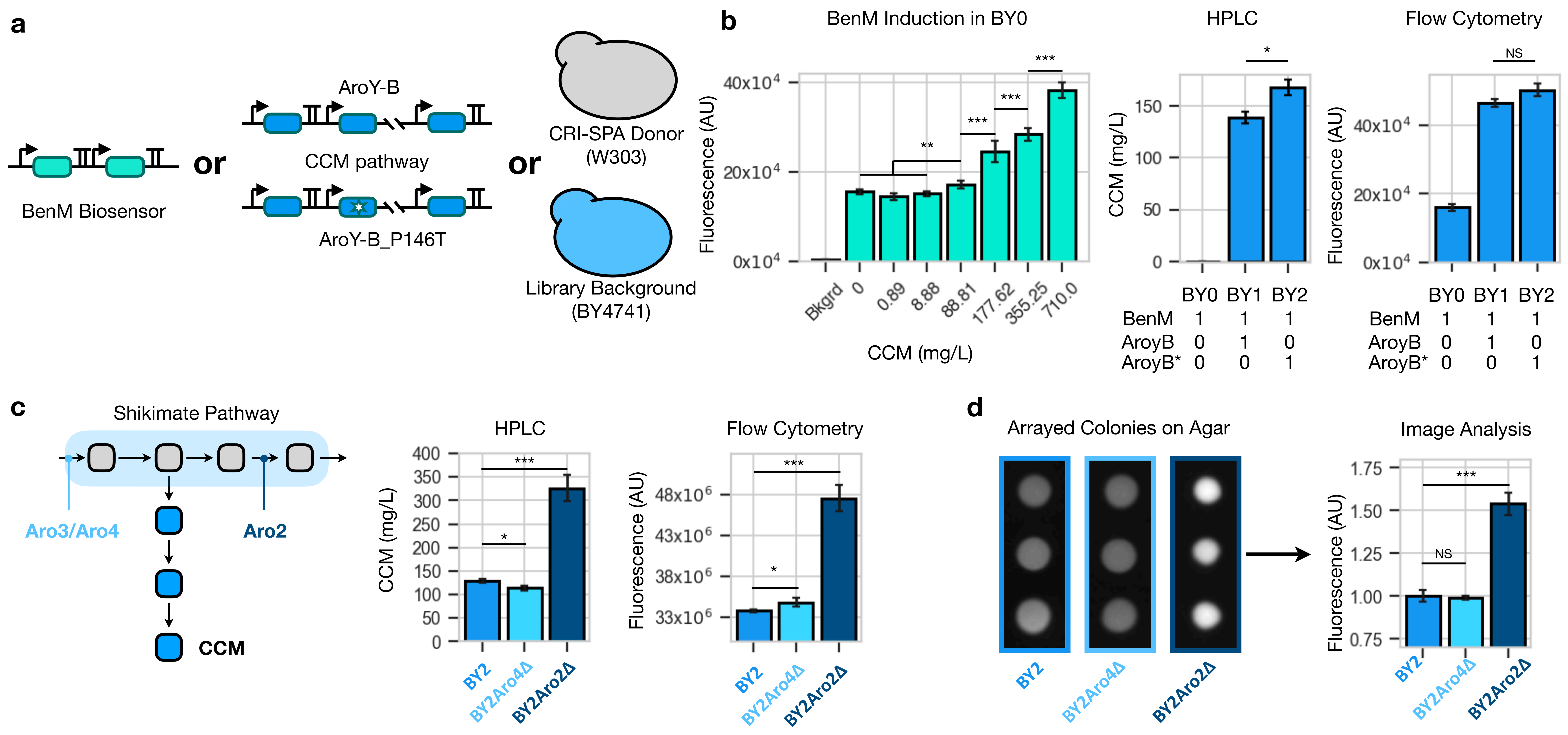


**Fig. 2. Characterization of CCM production and biosensing in CRI-SPA engineered yeast.**

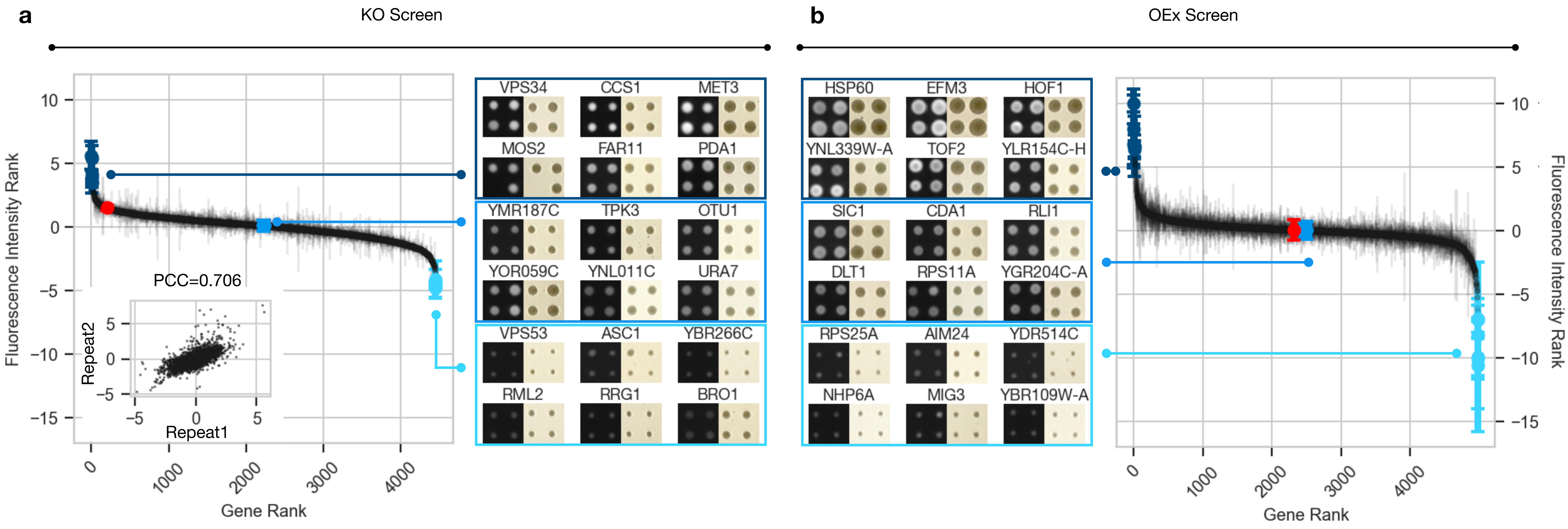


**Fig. 3. Ranking of host:pathway interactions for gene KO and overexpression.**

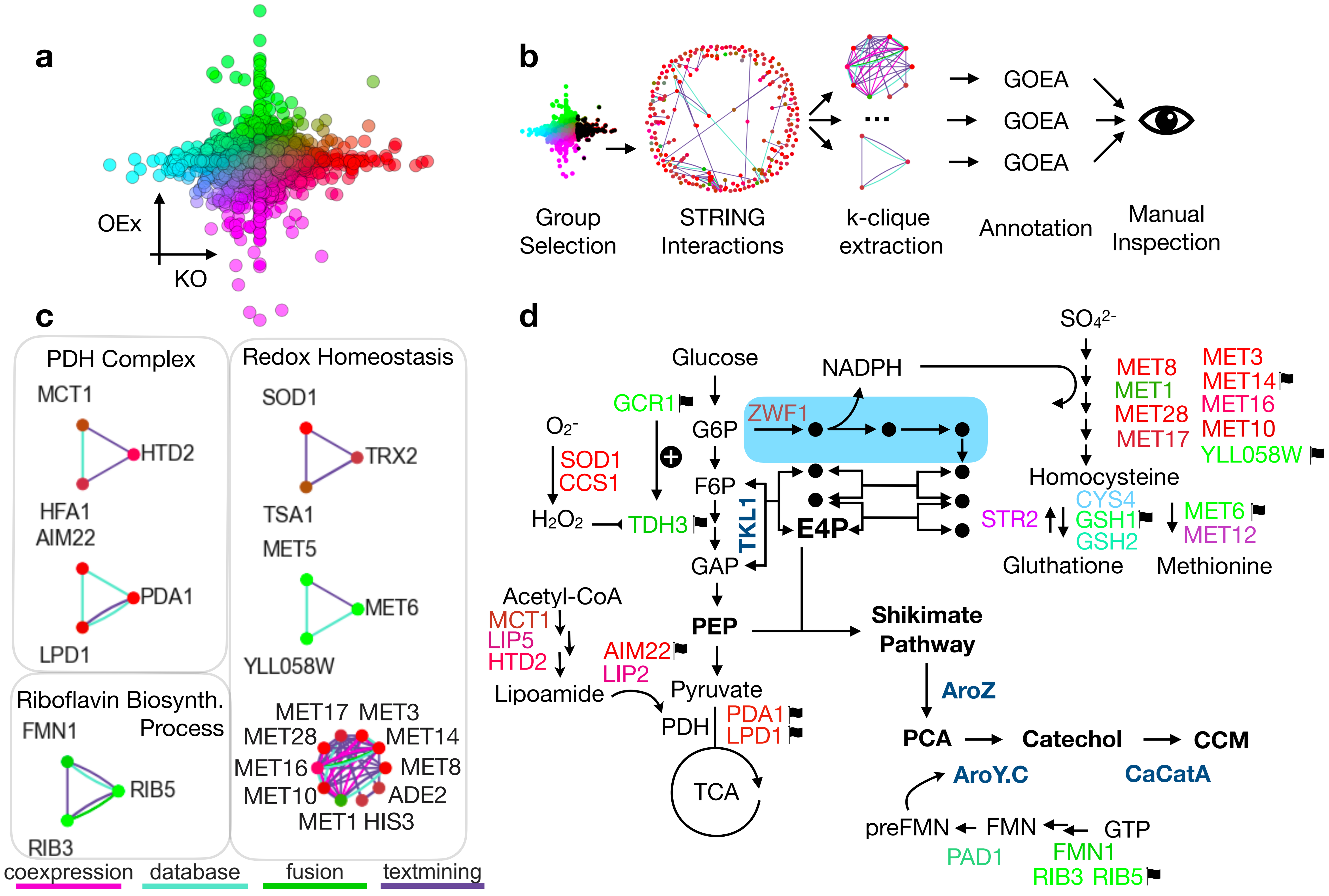


**Fig. 4. New cellular functions interacting with the synthesis of CCM.**

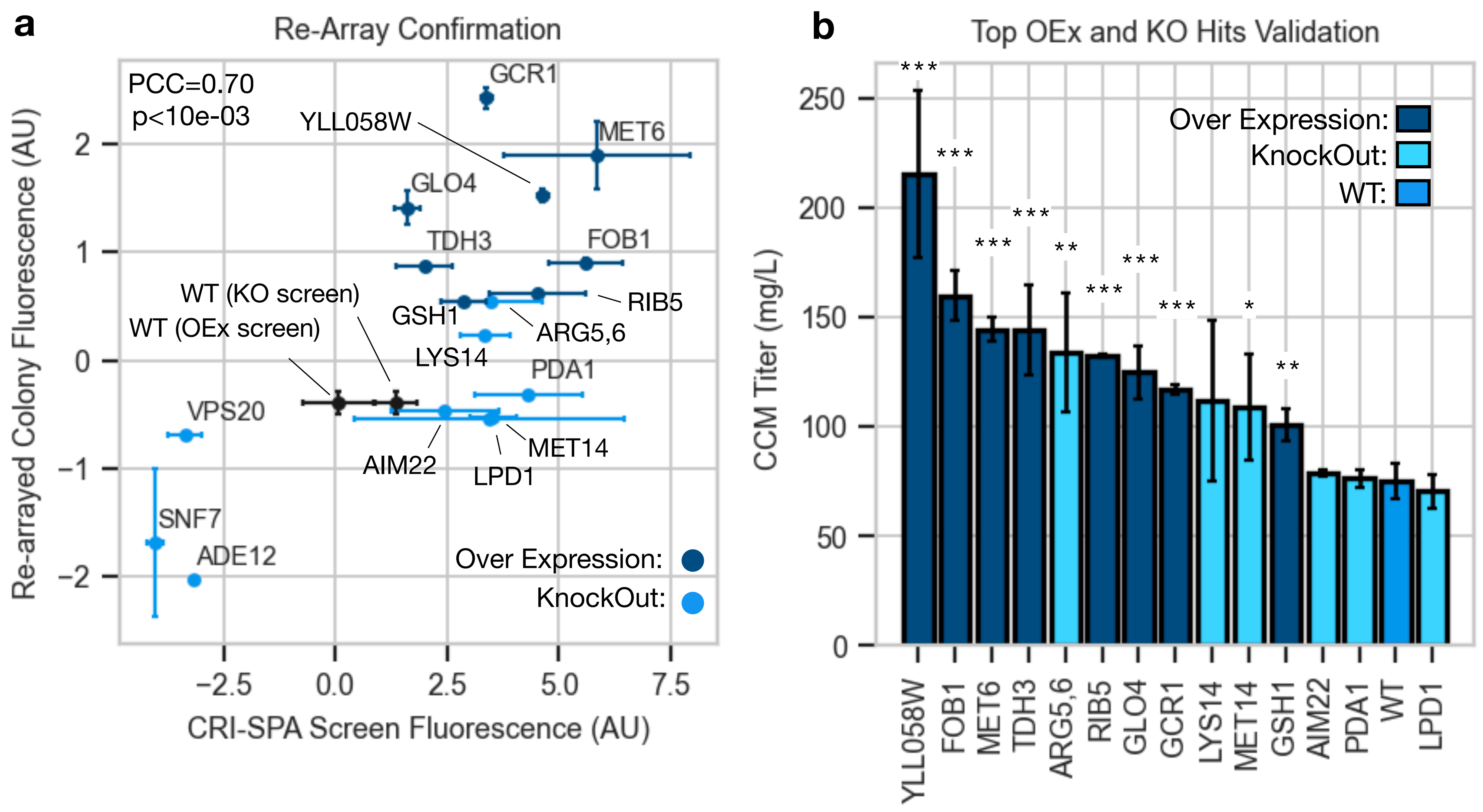


**Fig. 5. CRI-SPA hit validation.**

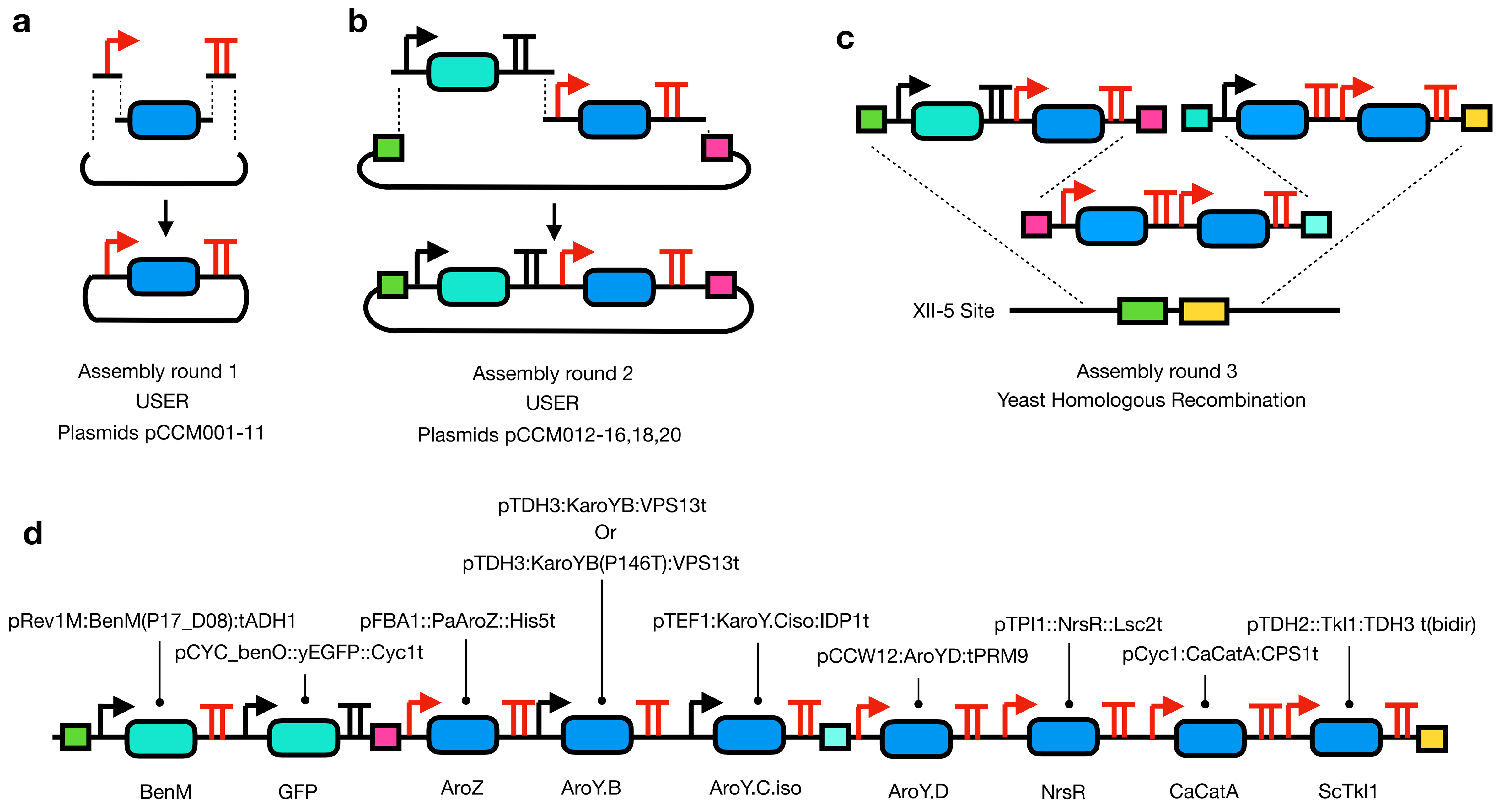


**Supplementary figure S1 Cloning of the CCM cassette.**

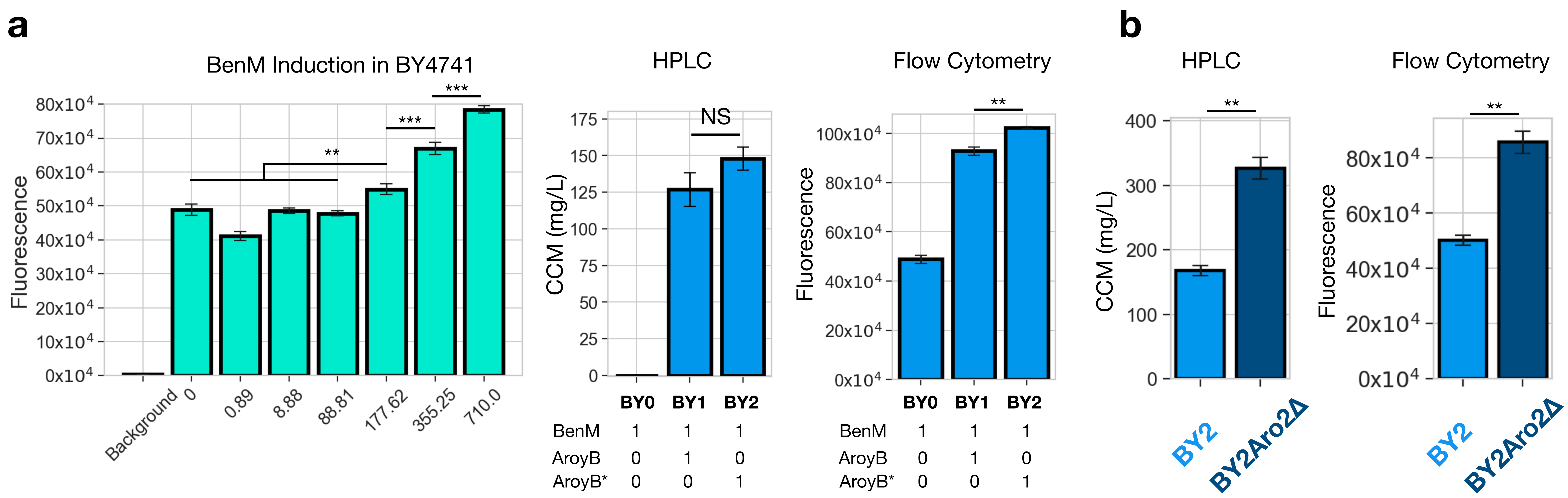


**Supplementary figure S2: Biological replicate of experiments in figure 2.**

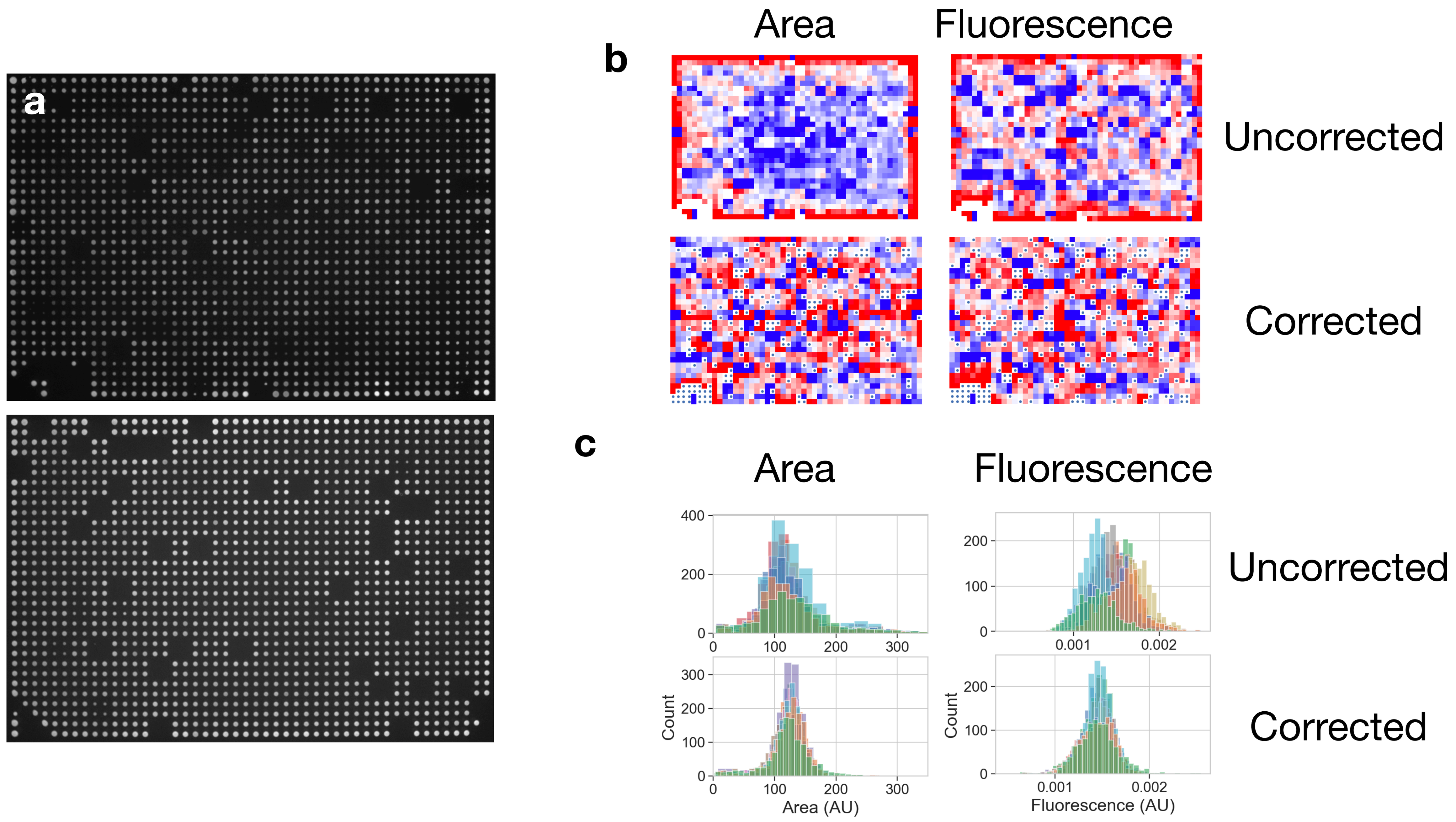


**Supplementary figure S3: Raw, uncorrected and corrected CRI-SPA data.**

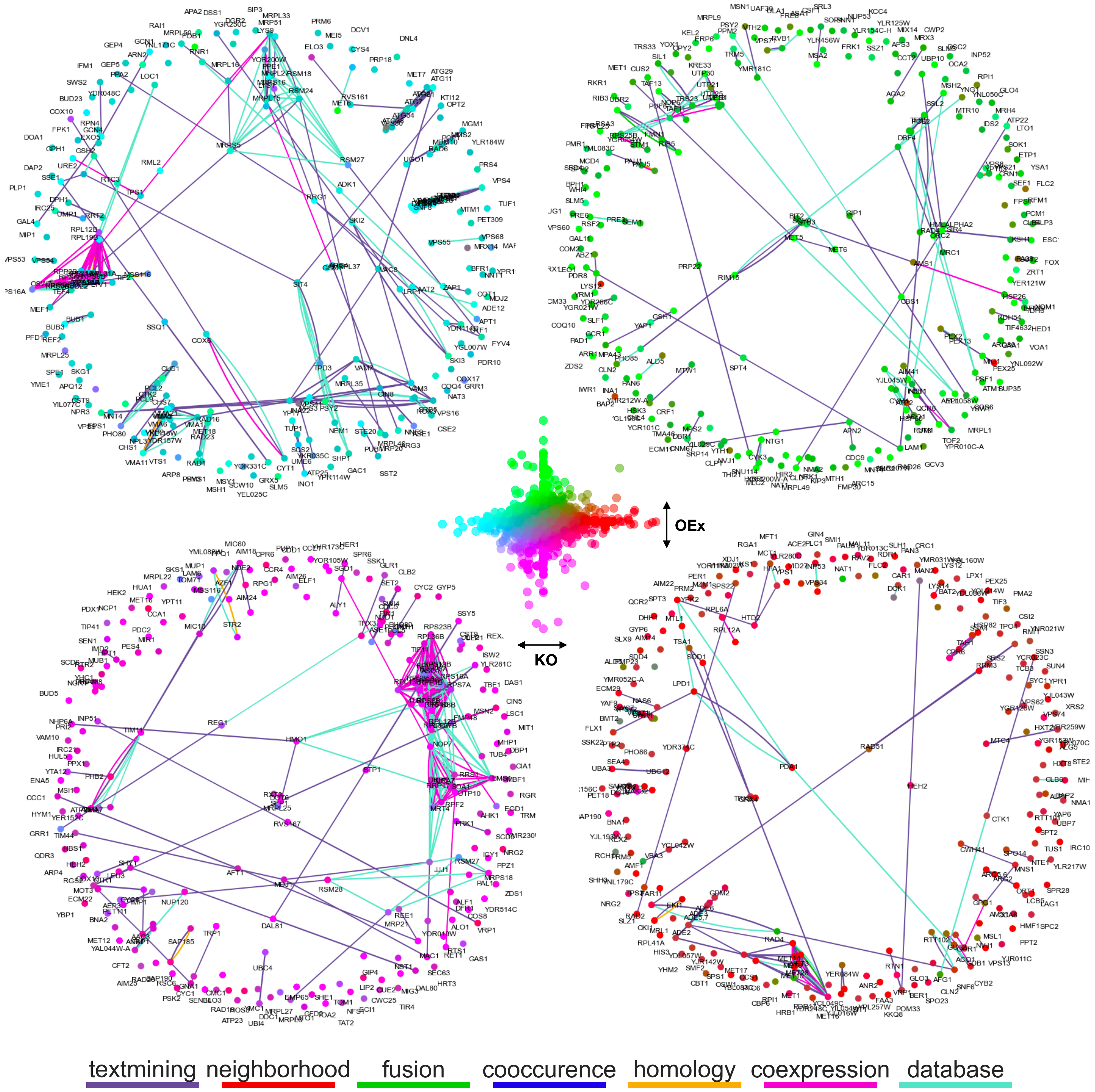


**Supplementary figure S4: STRING Clusters for 250 most extreme gene groups.**

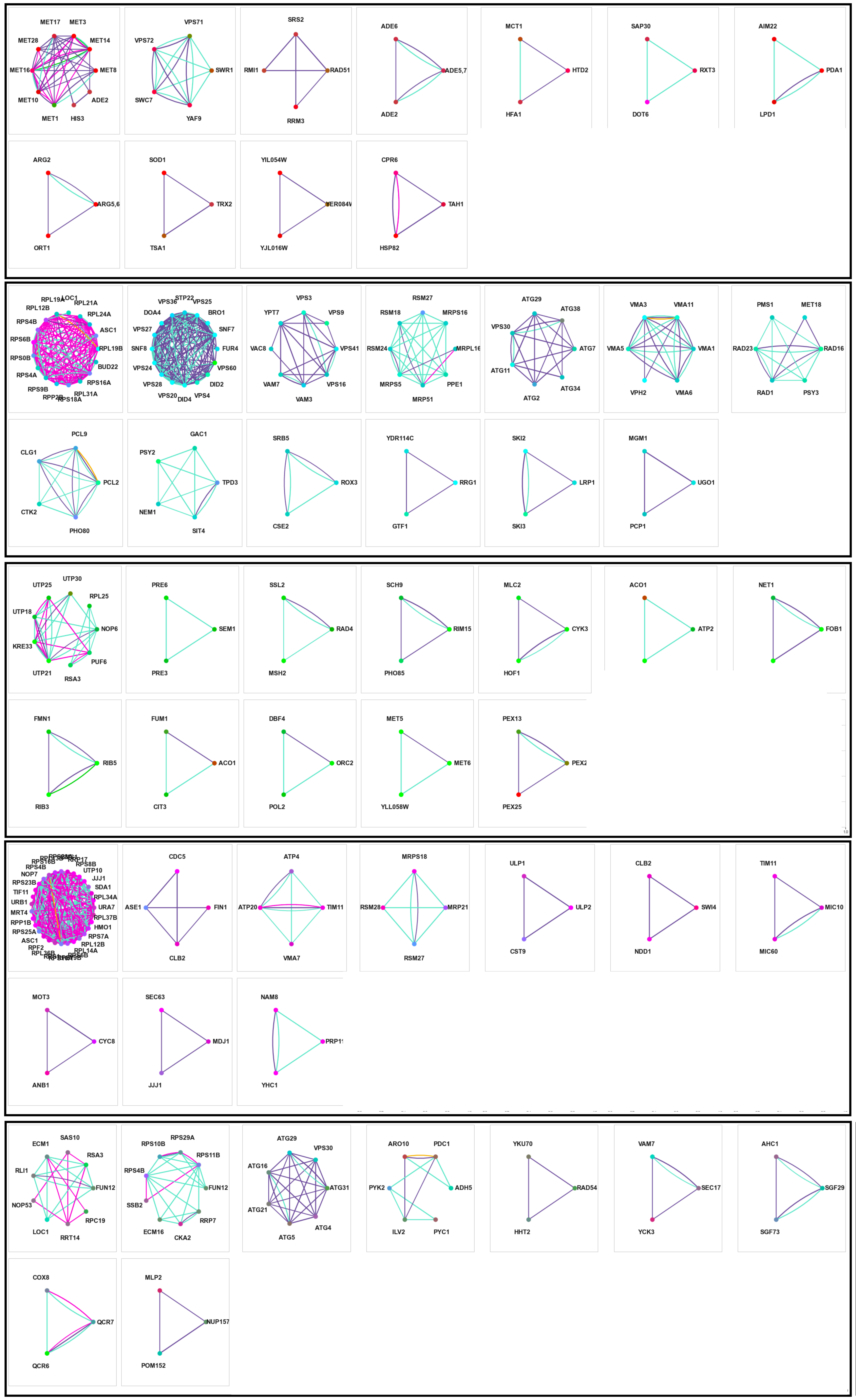

a

| | | | | | | |
|---|---|---|---|---|---|---|
| methionine biosynthetic process | chromatin remodeling | cellular response to DNA damage stimulus | purine nucleotide biosynthetic process | fatty acid Biosynthetic process | Rpd3L complex | Mitochond-rial pyruvate Dehydro-genase complex |
| arginine Biosynthetic process | antioxidant activity | None | protein folding | | | |

b

| | | | | | | |
|---|---|---|---|---|---|---|
| structural constituent of ribosome | endosome | fungal-type vacuole membrane | Mitochondrial small ribosomal subunit | autophagy | vacuolar acidification | DNA repair |
| cyclin-dependent protein kinase holoenzyme complex | protein phosphatase regulator activity | core mediator complex | None | RNA catabolic process | mitochondrial fusion | |

c

| | | | | | | |
|---|---|---|---|---|---|---|
| ribosome biogenesis | proteasome complex | DNA repair | protein kinase activity | cellular bud neck | Mitochon-drial nucleoid | nucleolus organization |
| riboflavin Biosynthetic process | Tricarbo-xylic acid cycle | mitotic DNA replication checkpoint signaling | None | peroxisomal membrane | | |

d

| | | | | | | |
|---|---|---|---|---|---|---|
| ribosome | spindle pole | proton trans-membrane transport | mitochondrial translation | SUMO-specific protease activity | None | Mitochond-rial crista junction |
| None | None | commitment complex | | | | |

e

| | | | | | | |
|---|---|---|---|---|---|---|
| ribosome biogenesis | Cytosolic small ribosomal subunit | phagophore assembly site | aromatic amino acid catabolic process to alcohol via Ehrlich pathway | None | fungal-type vacuole membrane | SAGA complex |
| respirasome | nuclear pore | | | | | |

**Supplementary figure S5: K-clique clusters extracted from STRING networks and annotated with GOEA.**

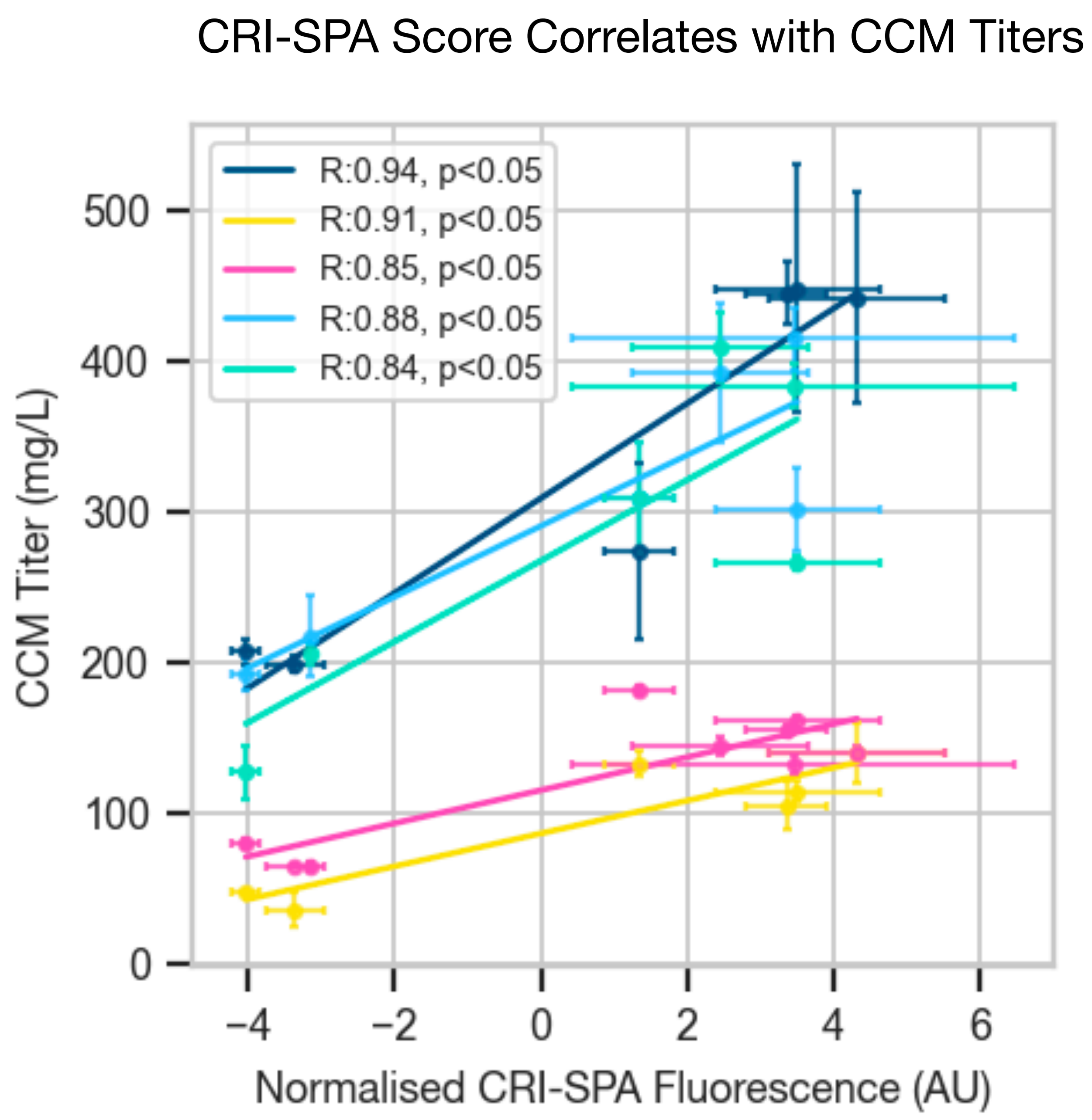


**Supplementary Figure S6: CRI-SPA fluorescence score correlates with CCM titers in liquid culture.**

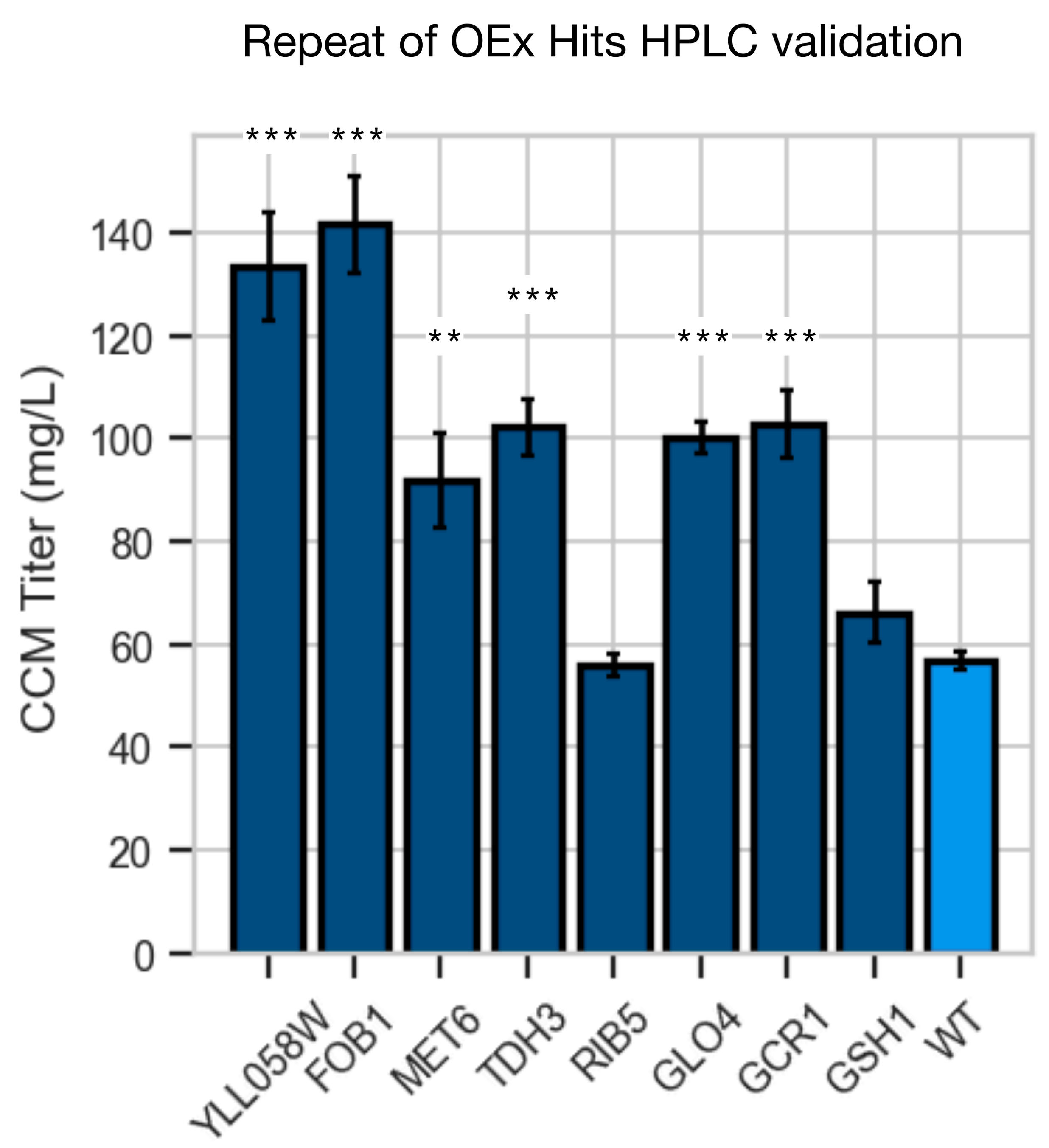


**Supplementary Figure S7: Biological replicate of experiment in figure 5.**
